# Differential Impact of rhamnolipids and TWEEN80 on composition and functionality of the gut microbiota in the SHIME system

**DOI:** 10.1101/2021.12.15.472660

**Authors:** Lisa Miclotte, Ellen De Paepe, Qiqiong Li, John Van Camp, Andreja Raikovic, Tom Van de Wiele

## Abstract

Dietary emulsifiers have been shown to affect the composition and function of the gut microbial community, both *in vivo* and *in vitro*. Yet, several knowledge gaps remain to be addressed: the impact from a longer timeframe exposure on the gut microbiota, interindividual variability in microbiome response and the putative impact from novel clean label alternatives for current food emulsifiers.

In the present study, the impact of one conventional dietary emulsifier, TWEEN80, and one potential novel alternative, rhamnolipids, on the human gut microbiota was investigated using the Mucosal Simulator of the Human Intestinal Microbial Ecosystem (M-SHIME). The faecal microbiota from two human donors, with high and low responsiveness to the emulsifiers, were exposed to 0,05 m% and 0,5 m% of the emulsifiers for 7 days.

The results confirmed previous observations that the effects on the composition and functionality are both emulsifier- and donor dependent. The effects reached an equilibrium after about 3 days of exposure. Overall, TWEEN80 and rhamnolipids displayed opposite effects: TWEEN80 increased cell counts, reduced propionate concentration, increased butyrate levels, increased a.o. *Faecalibacterium, Blautia* and *Hungatella* abundance, while rhamnolipids did the opposite. Rhamnolipids also sharply increased the abundance of unclassified Lachnospiraceae. On the other hand, both emulsifiers increased the relative abundance of unclassified Enterobacteriaceae. Both emulsifiers also altered the microbial metabolome in different ways and a pathway enrichment analysis tool revealed that the metabolome alterations could be reminiscent of gut issues and obesity.

Overall, the impact from the rhamnolipids was larger than that of TWEEN80 at similar concentrations, indicating that the former may not necessarily be a safer alternative for the latter. The microbiota’s response also depended on its original composition and the sensitivity status for which the faecal donors were selected, was preserved. Whether the same donor-diversity and longitudinal impact can be expected in the human colon as well and what impact this has on the host will have to be further investigated.

## 1. Introduction

Metabolic syndrome is defined as a set of conditions/symptoms that are risk factors for the development of the top most lethal diseases worldwide, notably cardiovascular disease (CVD) and diabetes [1]. Different definitions have been developed, each differing in their emphasis, but in general overweight or obesity and insulin resistance have been marked as the most important risk factors [1]. Western dietary patterns are strongly related to the development of metabolic syndrome. The overconsumption of saturated fat, sugar and salt and the general lack of dietary fiber in these diets have been evidenced to play a role in that relation. Western dietary patterns are often rich in (ultra-)processed food products, the consumption of which has recently been receiving more and more attention [2]. These highly processed food products often contain one or more food additives [3] of which emulsifiers and stabilizers are one large class. Several of these emulsifiers and stabilisers have recently been scrutinized for their purported destabilizing effects on the gut microbiota [4-6].

The human gut microbiota consists of a vast set of microorganisms that reside in the human gastro-intestinal tract [7,8]. It has been evidenced that it plays a role in or is influenced by obesity and metabolic aberrations in the host [9–12]. It is also known that nutrition has a profound and rapid effect on its composition and functionality [13,14].

TWEEN80, or polysorbate 80, is an emulsifier synthesized from fatty acids, sorbitol and ethylene oxide. It is authorized for use in food products like chewing gum, milk products, bakery wares, etc. by both EFSA and the FDA [15]. Its ADI has been set by EFSA at a level of 25 mg/kg bodyweight/day [16]. The impact of TWEEN80 on the gut microbiota has recently been investigated with respect to its putative impact on inflammatory bowel disease and weight gain [5,17]. In particular, Chassaing et al. (2015;2017) have found that TWEEN80 and carboxymethyl cellulose alter the composition of the gut microbiota and cause bacterial translocation across the intestinal mucus layer, where they are more prone to instigate inflammation in the gut epithelium.

Rhamnolipids are a type of glycolipid biosurfactants which are currently not applied in food products but which receive much attention because of their favourable emulsifying and eco-friendly properties [18–20]. They consist of a rhamnose or di-rhamnose backbone esterified to one or more fatty acids and are mostly produced by *Pseudomonas* species [21]. Because of their biotechnological production process, they would fit into the clean label movement, which stems from concern surrounding chemical additives and food processing and which aims amongst others to replace these chemicals by more natural alternatives. Whether rhamnolipids would also be safe with regards to intestinal health, is, however, unknown.

In our previous study, we investigated the impact of both TWEEN80 and rhamnolipids on the gut microbiota from 10 human donors *in vitro* and found the responses from the gut microbiota in terms of composition and functionality depended strongly on 1) the emulsifier and 2) the donor microbiota. TWEEN80 exhibited no significant effects on either microbiota composition, live-dead cell counts or the metagenome. Rhamnolipids, on the other hand, gravely decreased the living fraction of the bacterial population, increased propionate and decreased butyrate levels. Rhamnolipids also significantly altered the abundance of several microbial genera, among which a decrease in *Faecalibacterium, Bacteroides, Parabacteroides, Alistipes* and *Suterella* and an increase in *Escherichia/Shigella* and unclassified Enterobacteriaceae and *Phascolarctobacterium*.

The model in that study consisted of 48h batch incubations in a sugar-depleted SHIME-medium. The exposure to the emulsifier was thus rather short. In real life, consumers may be exposed to emulsifiers repeatedly, perhaps even on a daily basis. It is unknown how the gut microbiota reacts to such chronic emulsifier exposure in terms of composition and functionality. In the present study, the exposure time was therefore prolonged up to 7 days and the validated dynamic *in vitro* model known as the simulator of the human intestinal ecosystem or SHIME was utilized. From our preliminary screening of 10 human individuals, we selected the two donors with the most extreme responses and included LC-Orbitrap-MS based metabolomics analysis to gain a closer view on the impact of the emulsifiers on functionality. In this paper, we thus describe the effects of TWEEN80 and rhamnolipids on the composition and functionality of the gut microbiota from these two human donors. We also aim to compare the rhamnolipids to TWEEN80 in the context of the clean label movement.

## 2. Materials and methods

### 2.1 Experimental design

The impact of 2 dietary emulsifiers - TWEEN80 (P4780 - suitable for cell culture) and rhamnolipids (R90 - 90% pure, Sigma Aldrich) - on the human gut microbiota was investigated using the mucosal simulator of the human intestinal tract (M-SHIME^®^), a dynamic semi-continuous reactor-based model that mimics the different stages of the human gastro-intestinal tract (ProDigest-Ghent University). For this purpose, two independent M-SHIME experiments were executed, each inoculated with the faecal microbiota from a different human donor. In each experiment, two concentrations (0,05 m% and 0,5m%) of TWEEN80 and rhamnolipids were tested alongside a control condition, receiving water, in parallel sets of one stomach/small intestinal vessels and one colon vessel (Supplementary materials Figure 1). The mucosal environment was simulated through addition of mucin coated carriers in the colon compartments [22]. During the experiment, the double-walled vessels were continuously stirred by magnetic stirrers (Prosense) at 200 rpm and kept at 37°C through connection to a hot water bath (Julabo). The pH of the colon vessels was continuously monitored using pH electrodes (Consort) and controlled at 6,15 - 6,3 by semi-automatic addition of 0,5 M NaOH or 0,5 M HCl through pH controllers (Consort) and pumps (ProMinent).

### 2.2 Operation of the SHIME experiments

On the day of start-up, a faecal sample was harvested from a previously selected donor. The donors for these experiments were chosen from a pool of 10 individuals that were consulted for previous experiments [23]. Based on the response of the gut microbiota from these donors to the dietary emulsifiers in terms of SCFA-production, one high and one low responder was chosen. Both ended up being female, one 24 and the other 29 years old. Both had not received any antibiotic treatment in the three months prior to the faecal donation. The faecal samples were harvested in plastic lidded containers, which were rendered anoxic by addition of Anaerogen™ sachets (Oxoid Ltd.), and stored at 4°C for a maximum of 1 hour. A faecal slurry was then prepared as described in De Boever et al., (2000).

Before inoculation, the SHIME-model was operated on water for 24 hours. On the day of inoculation, all tubing and vessels were emptied, the stomach/small intestinal vessels were connected to pancreatic solution, containing 12,5 g/L NaHCO_3_ (Sigma Aldrich), 0,9 g/L porcine pancreatin (Sigma Aldrich) and 6 g/L Oxgall (BD), and nutritional SHIME feed (Prodigest PD – NM002A)(composition noted in Supplementary materials Table 1), acidified to a pH of 2,0. The colon vessels were filled with 360 mL of non-acidified nutritional SHIME medium, were then inoculated with 40 mL of faecal slurry, manually flushed with 100% N_2_ gas and left to acclimatize overnight.

The next day, the pumps were turned on and a first set of samples were taken, marking the official start of the experiment (0h). The feed – pumping scheme of the SHIME model is set out in Supplementary materials Table 2. Briefly, the stomach vessels were fed 140 mL of nutritional medium 3 times a day, as well as 60 mL of pancreatic juice, the feeding times 8 hours apart. From the stomach vessels, the solution was pumped to the colon vessels and after a residence time of 16h, discarded.

After at least 5 days of stable operating conditions, which was evaluated by means of pH-stability and visual inspection of the model, the microbial community was considered adapted to the SHIME-environment and the treatment was started. The pH was considered stable when it remained within a range of 6,15 –6,3 and visually, conditions were considered stable if the contents of the colon vessels remained turbid, indicating proper bacterial growth. Treatment solutions were prepared from rhamnolipids and TWEEN80 from Sigma Aldrich in distilled water and 10x more concentrated than the intended concentrations in the vessels. Three times a day, 40 mL of treatment solution was pumped into the respective stomach/small intestine vessels. After a treatment period of one week, the experiment was terminated.

To maintain anaerobic conditions during the experiments, all 14 SHIME-reactors were flushed for 5 min with N2 daily, between 9 and 10 am, except on days where mucin beads were replaced, in which case flushing occurred after bead replacement, around noon. Luminal samples were taken on days 2 (0h), 3, 5, 7, 8, 9, 10, 11 (twice: before and 1h after the start of the emulsifier treatment), 12, 13, 14, 16 and 18 (432h) for SHIME experiment 1 and days 2 (0h), 3, 5, 7, 8, 9, (twice: before and 1h after the start of the emulsifier treatment), 10, 11, 12, 14 and 16 (384h) for SHIME experiment 2. The full sampling scheme can be found in Supplementary materials Table 3 and 4. The procedure for taking luminal samples entailed attaching a 10 mL syringe to the sampling outlet on the lids of the colon reactors of the model, rinsing the syringe and the sampling tube of the colon vessel with the content of the vessel by filling and emptying the syringe multiple times, finally taking a 10 mL aliquot and dividing it in volumes of 1 mL over 1,5 mL Eppendorf tubes and one 10 mL tube for SCFA analysis.

Half of the 60 mucin beads (4 clusters of 15 beads) in each of the colon vessels were replaced every 2 days (except on weekends). First, mucin beads were coated in agar-mucin as described in Van den Abbeele et al., (2012). The beads were then sown together in clusters of 15 and stored in a sterilized container at 4°C until use. In the vessels, the beads were hanged inside a polyamide net, attached to lid with a nylon wire to facilitate exchange. Beads were always replaced in the morning between 10 and 12 am in the following way. First the lids of the vessels were connected with the N2 flush system and the vessels would be flushed to ascertain anaerobic conditions during the whole procedure. Next, the vessels were opened one by one, 2 of the 4 clusters in the vessels were replaced by 2 fresh ones, the vessels were closed and flushed with N_2_-gas for a further 5 min. Meanwhile the beads were centrifuged for 3 minutes at 500 *g* to collect the mucin.

### 2.3 SCFA analysis

The SCFA-concentrations were determined by means of diethylether extraction and capillary gas chromatography coupled to a flame ionization detector [25,26]. Briefly, 1 mL aliquots were were diluted 2x with 1 mL mili-Q water and SCFA were extracted by adding approximately 400 mg NaCl, 0,5 mL concentrated H_2_SO_4_, 400 μL of 2-methyl hexanoid acid internal standard and 2 mL of diethyl ether before mixing for 2 min in a rotator and centrifuging at 3000 *g* for 3 minutes. Upper layers were collected and analysed using a GC-2014 capillary gas chromatograph (Shimadzu), equipped with a capillary fatty acid-free EC-1000 Econo-Cap column (Alltech), 25 m × 0,53 mm; film thickness 1,2 μm, and coupled to a flame ionization detector and split injector. The resulting SCFA-levels were visualized in R (version 4.1.0) by use of lineplots created using ggplot2 (v3.3.3).

### 2.4 Total cell counts

The impact of the emulsifiers on total cell concentrations was assessed by first performing SYBR^®^ green staining, on luminal SHIME-samples after which cells were counted on an Accuri C6+ flow cytometer from BDbiosciences Europe. Samples were analyzed every few days in batches from frozen at −20°C. Dilutions up to 10 000x were prepared in 96-well plates using 0,22 μm filtered sterile 0,01 M phosphate buffered saline (PBS) (HPO_4_^2-^ / H_2_PO_4_^-^, 0,0027 M KCl and 0,137 M NaCl, pH 7,4, at 25 °C) and these were subsequently stained with SYBR^®^ green (SG) (100x concentrate SYBR^®^ Green I, Invitrogen, in 0,22 μm-filtered dimethyl sulfoxide) [27,28]. After incubation for 25 minutes, the total populations were measured immediately with the flow cytometer, which was equipped with four fluorescence detectors (530/30 nm, 585/40 nm, >670 nm and 675/25 nm), two scatter detectors and a 20 mW 488 nm laser. The flow cytometer was operated with Milli-Q (MerckMillipore) as sheath fluid. The blue laser (488 nm) was used for the excitation of the stains and a minimum of 10 000 cells per sample were measured for accurate quantification. Settings used were an FLH-1 limit of 1000, a measurement volume of 25 μL and the measurement speed was set to ‘fast’. Cell counts were obtained by gating the total cell populations in R (version 4.1.0) according to the Phenoflow-package (v1.1.6)[29]. Gates were verified using data from cell-free control samples (0,22 μm filter sterilized 0,01M PBS) (Supplementary Figure 1). Lineplots of total cell counts were created using ggplot2 (v3.3.3).

### 2.5 Amplicon sequencing

Luminal samples from day 2, 11, 12, 14, 18 from SHIME 1 and day 2, 9, 10, 12 and 16 from SHIME 2 and mucosal samples from day 4, 11, 14, 16 and 18 from SHIME 1 and day 4, 9, 11, 14 and 16 from SHIME 2 were selected for Illumina 16S rRNA gene amplicon sequencing. Extraction and quality verification of DNA, library preparation and 16S rRNA gene amplicon sequencing were performed as described in Miclotte 2020.

Recovery of OTU’s from the amplicon data was carried out using mothur software version 1.40.5 and guidelines [30]. First, contigs were assembled, resulting in 9 815 109 sequences, and ambiguous base calls were removed. Sequences with a length of 291 or 292 nucleotides were then aligned to the silva_seed nr.123 database, trimmed between positions 2 and 13 423 [31]. After removing the sequences containing homopolymers longer than nine base pairs, 2 222 311 (99%) of the unique sequences were retained. A pre-clustering step was then performed, allowing only three differences between sequences clustered together and chimera.vsearch was used to remove chimeras, retaining 73% of the sequences. The sequences were then classified using a naïve Bayesian classifier against the Ribosomal Database Project (RDP) 16S rRNA gene training set version 16, with a cut-off of 85% for the pseudobootstrap confidence score. Sequences that were classified as Archaea, Eukaryota, Chloroplasts, unknown, or Mitochondria at the kingdom level were removed. Finally, sequences were split at the order level into taxonomic groups using the OptiClust method with a cut-off of 0.03. The data were classified at a 3% dissimilarity level into OTUs, resulting in a .shared (count table) and a .tax file (taxonomic classification).

For the entire dataset of 140 samples, 137 458 OTUs were detected in 199 genera. An OTU was here defined as a collection of sequences with a length between 291 and 292 nucleotides and with 97% or more similarity to each other in the V4 region of their 16S rRNA gene after applying hierarchical clustering.

#### Bioinformatics in R

The shared and taxonomy files resulting from the mothur pipeline were loaded into R for further processing. Absolute singletons (OTUs with only one read over all samples) were removed, resulting in 28 128 OTUs being retained [32]. Rarefaction curves were created to evaluate the sequencing depth (Supplementary Figure 2) [33]. Relative and absolute abundances of the OTUs and genera were calculated from the read counts and were explored via bar plots using ggplot2 (v3.3.3) and via principle coordinate analysis (PCoA) on the abundance based Jaccard distance matrix using the prcomp-function in the stats (v4.1.0) package. Relative abundances were calculated as percentages of the total read counts per sample. Absolute abundances (cfr. quantitative microbial profiling) were calculated by multiplying the total cell counts obtained via flow cytometry with the relative abundances of the OTUs (similar to Vandeputte et al., (2017)).

The Chao1, Shannon and Simpson diversity indices were calculated for the microbial community based on the OTU-table using the SPECIES (v1.0) package and the diversity function in the vegan (v2.5-7) package. Indices were plotted using ggplot2.

To investigate the effects of the individual constraints on the microbial community, a series of distance based redundancy analyses (dbRDAs) was then performed on the scores obtained in the PCoA on the Jaccard distance matrix using the capscale function in the vegan (v2.5-7) package. Permutation tests were used to evaluate the significance of the models and of the explanatory variables (De Paepe et al., 2018). DbRDAs were created for both the relative and absolute abundance dataset.The global model included the factors Treatment (a concatenation of the factors Emulsifier and Emulsifier concentration) Timepoint, and Donor as explanatory variables and the absolute abundances of the genera as explanatory variables. In three subsequent dbRDAs, the constrained variance attributed to each of the factors was investigated by each time conditioning out all but one factor. The results of the dbRDAs were plotted as Type II scaling correlation triplots showing the two first constrained canonical axes (labeled as dbRDA Dim 1/2) and the proportional constrained eigenvalues representing their contribution to the total (both constrained and unconstrained) variance. The sites were calculated as weighed sums of the scores of the response variables.

Finally, to evaluate which genera were likely differentially abundant between emulsifier treatments and control, the DESeq2 package (v 1.32.0) was applied on the read count-table at genus level. To this end, only the read counts from the samples in the treatment period were used.

To streamline the DESeq-process, pre-filtering according to McMurdie & Holmes, (2014) was first applied on the read count-table, after which a genus-level table was created using the aggregate function (stats package v4.1.0). In the generalized linear model, the factor Timepoint, Donor, and Treatment were included. A likelihood ratio test was employed within the DESeq function on the reduced model, containing only the factors Donor and Timepoint, to test for the significance of the model. Low count genera were subjected to an empirical Bayesian correction [35]. For pairwise comparison of TWEEN80 and rhamnolipid treatments versus controls, Wald tests were used after shrinkage of the Log2FoldChange (L2FC) values by means of the lfcShrink function with the adaptive shrinkage estimator from the ashr-package [36]. P-values were adjusted by means of a Benjamini-Hochberg procedure [35]. Results were visualized in volcanoplots, displaying the -log(adjusted p-value) versus the Log2FoldChange of each genus. Additionally, box plots were created showing the log-transformed pseudocounts extracted by the plotCounts function for each genus that showed significant differential abundance.

### 2.6 Metabolomics

Both targeted and untargeted polar metabolomics analyses of luminal SHIME-samples from days 2, 11, 12, 14 and 18 from SHIME 1 and days 2, 9, 10, 12, 16 from SHIME 2 were performed in collaboration with the Lynn Vanhaecke lab according to previously described protocols [37–39].

#### Polar metabolomics analysis

Polar metabolites were first extracted from the frozen SHIME-samples by means of ultra pure water (0,055 μS cm^-1^) obtained via a purified water system (VWR International, Merck). For this purpose, 1,5 mL SHIME-sample was first spiked with 75 μL internal standard solution (100 ng/μL L-alanine-d3 and D-valine-d8) and then centrifuged for 5 min at 13 300*g*. The supernatant was then filtered over a 0,22 μm polyvinylidene difluoride filter (Millipore), diluted 1:5 using ultrapure water and transferred into an HPLC-vial [37].

Polar metabolites were then separated and detected using an ultra-high performance liquid chromatograph system coupled to a Q-Exactive™ mass spectrometer with heated electronspray ionisation (UHPLC-HRMS Thermo Fisher Scientific) according to validated methods as described in De Paepe et al 2018 and De Spiegeleer et al 2020. Separation was achieved using a Dionex Ultimate XRS 3000 system (Dionex) equipped with a Acquity UPLC HSS T3 column (1,8 μm particle size, 2,1 mm internal diameter, 150 mm length, Waters) kept at a constant 45°C during analysis. Samples were injected in randomized order at a volume of 10 μL and eluted using a polar to apolar gradient formed using Ultrapure water (A) and LC/MS grade acetonitrile (Thermo Fisher Scientific) (B), both acidified with 0,1% LC/MS grade formic acid (Thermo Fisher Scientific). Elution occurred at a constant flow rate of 0,4 mL/min. The gradient profile comprised the following proportions of solvent A: program: 0 – 1,5 min at 98%, 1,5 – 7,0 min from 98% to 75%, 7,0 – 8,0 min from 75% to 40%, 8,0 – 12,0 min from 40% to 5%; 12,0 – 14,0 min at 5%, 14,0 – 14,1 min from 5 to 98%, followed by 4,0 min of re-equilibration [37,38].

Subsequent detection of the metabolites occurred using a Q-exactive™ Orbitrap mass spectrometer (Thermo Fisher Scientific), with a heated electrospray ionization source (HESI – II) operated in polarity switching mode (0/B/1 position). The system was operated under the following settings: m/z range of 53,4 - 800 Da, a maximum injection time of 70 ms, a mass resolution of 140 000 full width at half maximum and an automatic gain control of 10^6^ ions. The sheath, auxiliary and sweep gas (N_2_) flow rate for the HESI-II source were set at 50, 25 and 3 arbitrary units. The heater and capillary temperatures were set at 350 °C and 250 °C respectively, the S-lens RF-level was 50 V and the spray voltage was 4 kV [39].

Before analysis of the samples, the both ionization modes of the Q-Exactive system were calibrated using ready-to-use calibration mixtures (Thermo Fisher Scientific) for optimal mass accuracy. Sample analysis was also preceded by injection of a standard mixture followed by injection of a series of four quality control samples – obtained by pooling equal aliquots from all samples together – to allow system stabilization. Duplicate quality control samples were also added intermittently between series of 10 samples, to allow for correction of instrument instability and signal drift.

#### Data Processing

##### Targeted metabolomics

After data acquisition, peak identification and quantification was performed within Xcalibur 3.0 software (Thermo Fisher Scientific). Metabolite peaks were manually identified using an in-house retention time library, the peak location in the standard mixture and the C13/C12 ratio according to CD 2002/657/EC guidelines. A minimal peak intensity threshold of 500 000 a.u. was employed. Peak areas were gathered as one table in Excel 2016 and normalized through division by peak area values for the internal standard (D-valine-d3). This base table containing area ratio’s was then plugged into R (version 4.1.0) for further principle coordinate analysis or split up per treatment versus the control condition for analysis in MetaboAnalyst 5.0 (2021).

Disease signatures present in the metabolite dataset were explored using the quantitative Enrichment Analysis application in MetaboAnalyst 5.0 (Xia Lab, McGill University). Prior to these analyses, log transformation and Pareto scaling were performed via the website tool. The faecal library was chosen as reference database. The Statistical Analysis application on MetaboAnalyst was also employed to identify differential compounds. Compounds were considered differential when they were indicated as significantly different by the Wilcoxon Rank-sum test, the VIP value was over 1 in PLS-DA and if p(1) > 0,1 and p(corr)[1] > 0,4 in OPLS-DA for treatment with both concentrations with TWEEN80 and rhamnolipids.

##### Untargeted metabolomics

Compound Discoverer 3.0 (Thermo Fisher Scientific) was used first for automated peak extraction, peak alignment, deconvolution and noise removal with settings as given in Table 1 [38,39]. Suspect compounds were identified based on the Chemspider database (Royal Society of Chemistry).

**Table 1:** Settings utilized in Compound Discoverer for peak extraction and compound detection.

| Settings in Compound Discoverer 3.0 | Value |
| --- | --- |
| Peak intensity threshold | 500 000 a.u. |
| Specific adducts | [M+H], [M+K],[M+Na], [M-H], [M-K], [M-Na] |
| Retention Time | 0,5 - 16 |
| S/N threshold | 15 |
| Max peak shift | 0,2 min |
| Mass Tolerance | <5 ppm |
| Max peak width | 0,55 |

**Table 1:** P-values of Wilcoxon rank sum tests (with Holm correction) comparing SCFA-value differences of treatment vessels with controls before and after the start of the treatment.

|  | Donor | Acetate | Propionate | Butyrate | Total |
| --- | --- | --- | --- | --- | --- |
| **P80 – 0,05%** | 1 | 0,11 | 0,34 | 0,69 | 0,49 |
|  | 2 | **0,03** | **0,06** | **0,06** | 0,34 |
| **P80 – 0,5%** | 1 | 0,20 | **0,06** | 0,11 | 0,49 |
|  | 2 | 0,69 | **0,03** | **0,06** | 0,49 |
| **RL – 0,05%** | 1 | 0,89 | 0,69 | **0,03** | 0,20 |
|  | 2 | 0,20 | **0,03** | **0,03** | 0,34 |
| **RL – 0,5%** | 1 | 0,69 | **0,06** | 0,34 | 0,69 |
|  | 2 | 0,69 | 0,34 | **0,03** | 0,20 |

Peak area data obtained from Compound Discoverer was plugged into SIMCA 17 (Umetrics) for multivariate statistical analysis. To this end, first QC-normalisation and QC-filtering based on a CV < 30% were performed to eliminate instrumental signal drift and metabolites for which detection was unstable. QC-normalisation was performed per compound by dividing the peak area in a sample by the means of the interspersed duplicate QC-samples that follow the batch of 10 samples that sample belongs to.

In SIMCA, a PCA was first generated as an exploration of overall data patterns. Next, Orthogonal Partial Least Squares Discriminant Analysis (OPLS-DA) was used generate predictive models comparing EM-treatment conditions to the emulsifier free control condition. For each of these comparisons a model with a Q2 > 0,5, R2Y ≈ 1 and R2X ≈ 1 were selected, of which the significance was verified by ANOVA and Permutation tests. Model creation involved compound-filtering based on the Variable importance of the Projection criterion: VIP >1. Compounds characteristic for the particular EM-treatment were then identified based on their VIP (>1), a positive Jackknife confidence interval and the S-plots (|p(1))| > 0,05 and relevance |p(corr(1))| > 0,4).

## 3. Results

### 3.1 Quality control

During the SHIME experiments, the performance of the model was evaluated by measuring and adjusting the pH in the vessels as well as following up the produced SCFA-levels and cell concentrations. These concentrations were all at normal levels, from which we concluded that the research questions can be answered.

### 3.2 Cell counts

TWEEN80 mildly increased the total cell counts compared to the control, while 0,5 m% rhamnolipids decreased them, in a concentration dependent manner. Near the end of the treatment period, the cell counts from the compartments treated with rhamnolipids recovered slightly. High and low responder status of the selected microbiota weren’t particularly visible here (Figure 1).

**Figure 1:**
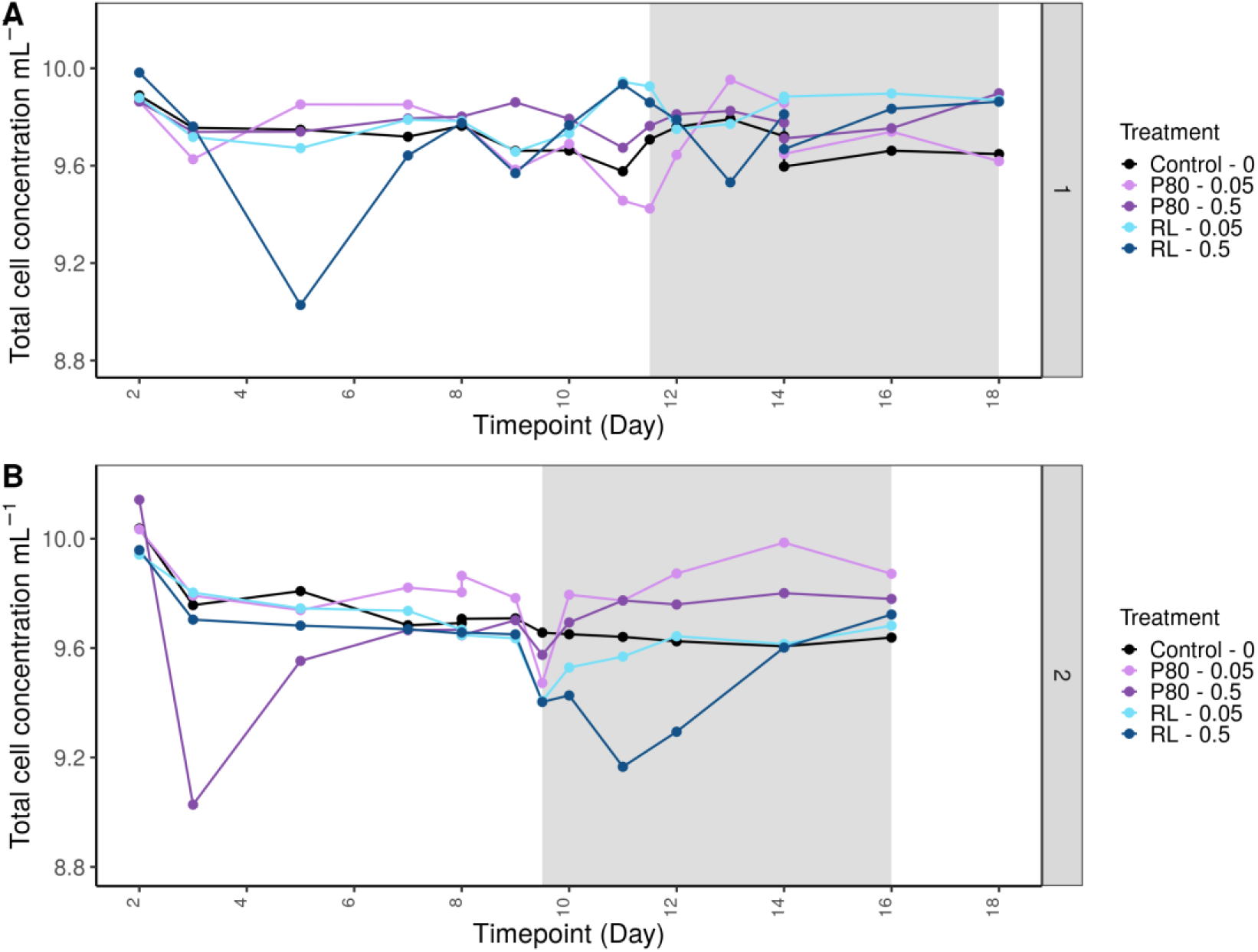
Impact of emulsifiers on total cell concentration. Total cell concentration (cells/mL) was measured in luminal suspension during a 16 and 18 day SHIME experiment investigating the impact of TWEEN80 and rhamnolipids (0,05 m% and 0,5 m%) on the gut microbiota of two human faecal donors. Donor 1 was selected for low emulsifier sensitivity and donor 2 for high emulsifier sensitivity. The 7-day treatment period is indicated by the grey background.

### 3.3 Amplicon sequencing

The impact of emulsifier treatment on microbial community composition was investigated using 16S rRNA amplicon sequencing on an Illumina Miseq platform. This analysis revealed that both emulsifiers altered the microbial composition, and that the response depended on the initial microbial composition (i.e. the donor), the emulsifier and the intestinal environment (mucus *vs* lumen).

First, a general assessment of trends within the microbial communities was performed using PCoA and dbRDA on both the absolute and relative abundance data. The dbRDA’s revealed that the factors Timepoint, Donor and Environment explained the largest share of the total constrained variance in the data. For the relative abundance data Timepoint, Donor and Environment explained respectively 12,94%, 7,18% and 5,67% and for the absolute abudances the factors Timepoint and Donor accounted for 17,26% and 8,08% (Figure 2; Supplementary Figure 10). The factor Treatment, a concatenation of the applied emulsifier and its concentration represented only 8,31% of the variance in the absolute abundance data and 5,49% of the variation in the relative abundance data (Supplementary Figure 10). The effects of all of the factors were significant (p < 0,005).

**Figure 2:**
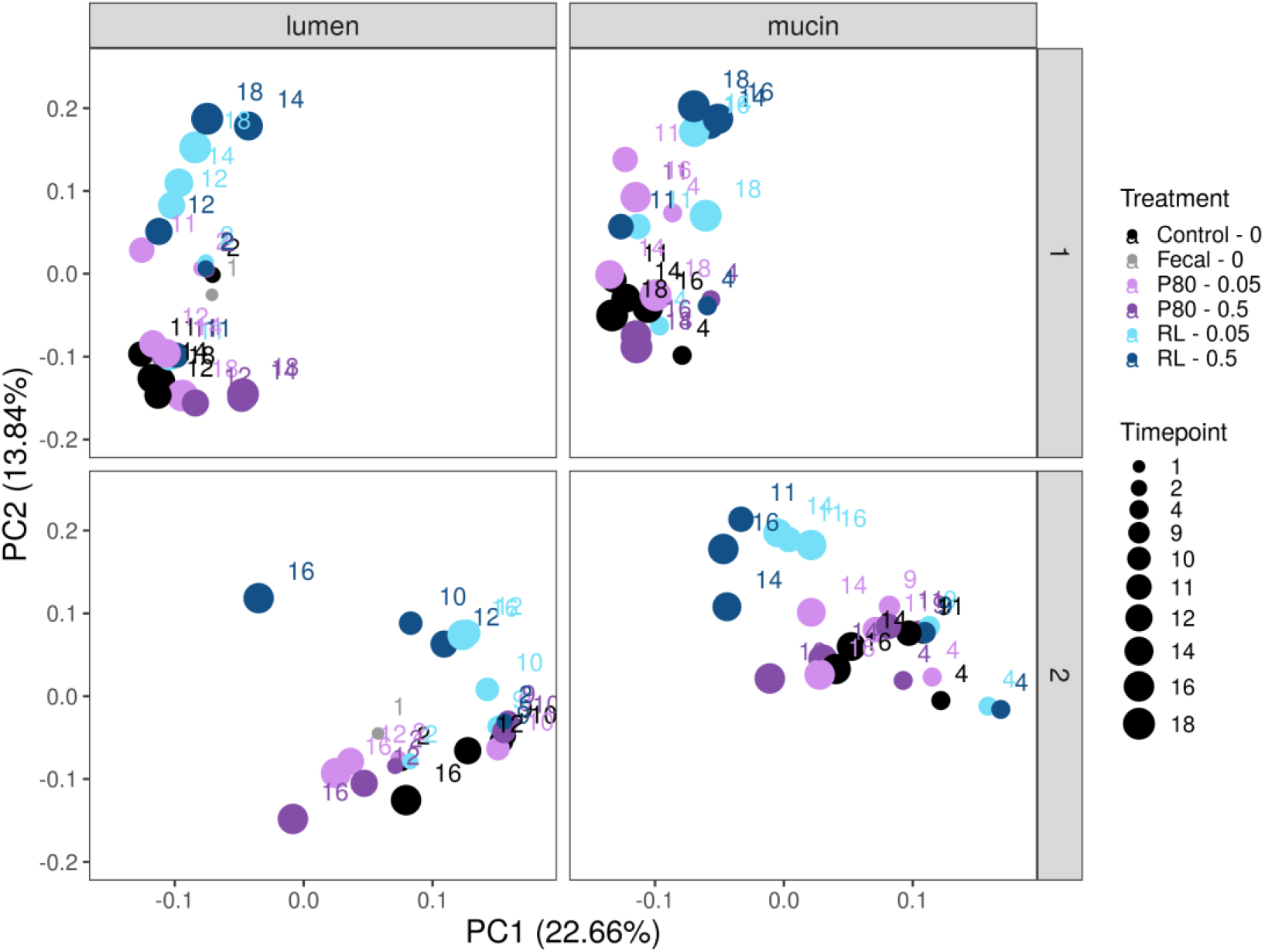
Principle coordinate analysis (PCoA) of 16S rRNA Illumina amplicon sequencing data extracted from luminal suspensions collected during a 16 and 18 day SHIME experiment investigating the impact of TWEEN80 and rhamnolipids (0,05 m% and 0,5 m%) on the gut microbiota from two human faecal donors. Donor 1 was selected for low emulsifier sensitivity and donor 2 for high emulsifier sensitivity. For each condition, different timepoints are indicated relative to the start of the treatment: timepoint 0 (in days).

Although the effect of the emulsifier treatment differed considerably between donors and the intestinal niches, both emulsifiers caused substantial alterations to the structure of the microbial communities. The treatment with TWEEN80 resulted in increased absolute abundances of *Faecalibacterium* and *Blautia* in donor 1 and also *Agathobacter* and *Subdoligranulum* in donor 2 (Figure 3&4). The rhamnolipids, especially at the highest concentration, strongly reduced the absolute abundance of all genera, except the unclassified Lachnospiraceae in both donors and *Megamonas* in donor 2 (Figure 3&4). Investigation of the relative abundances of both luminal and mucosal compartments revealed that in the mucosal compartment, the presence of unclassified Lachnospiraceae was more pronounced (Supplementary Figures 2 & 3).

**Figure 3:**
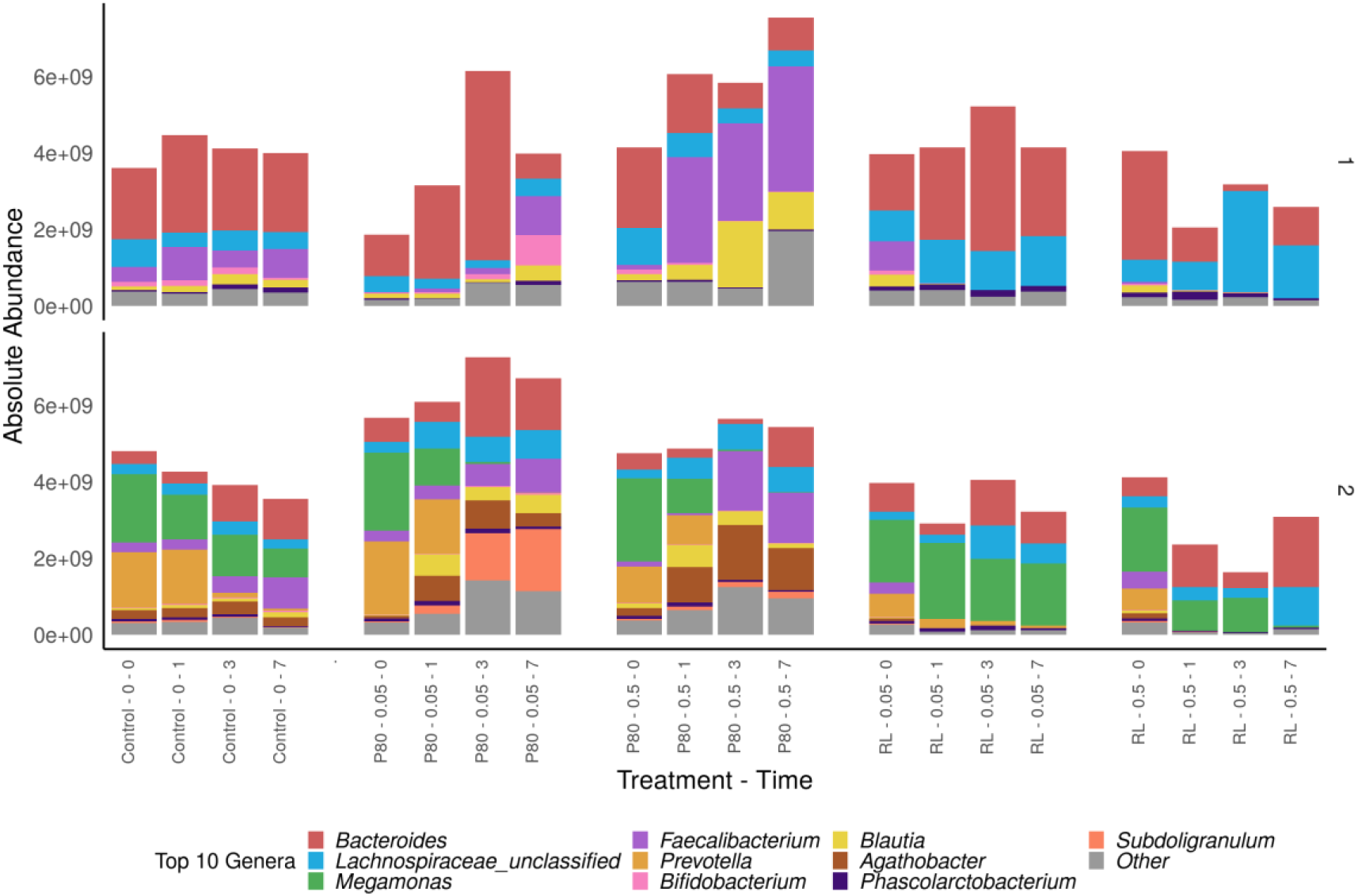
Absolute abundance (in cells/mL) of 10 most abundant genera derived from 16S rRNA gene amplicon sequencing, measured in luminal suspensions of a 16 and 18 day SHIME experiment investigating the impact of TWEEN80 and rhamnolipids (0,05 m% and 0,5 m%) on the gut microbiota from two human faecal donors. Donor 1 was selected for low emulsifier sensitivity and donor 2 for high emulsifier sensitivity. For each condition, different timepoints are indicated relative to the start of the treatment: timepoint 0 (in days).

**Figure 4:**
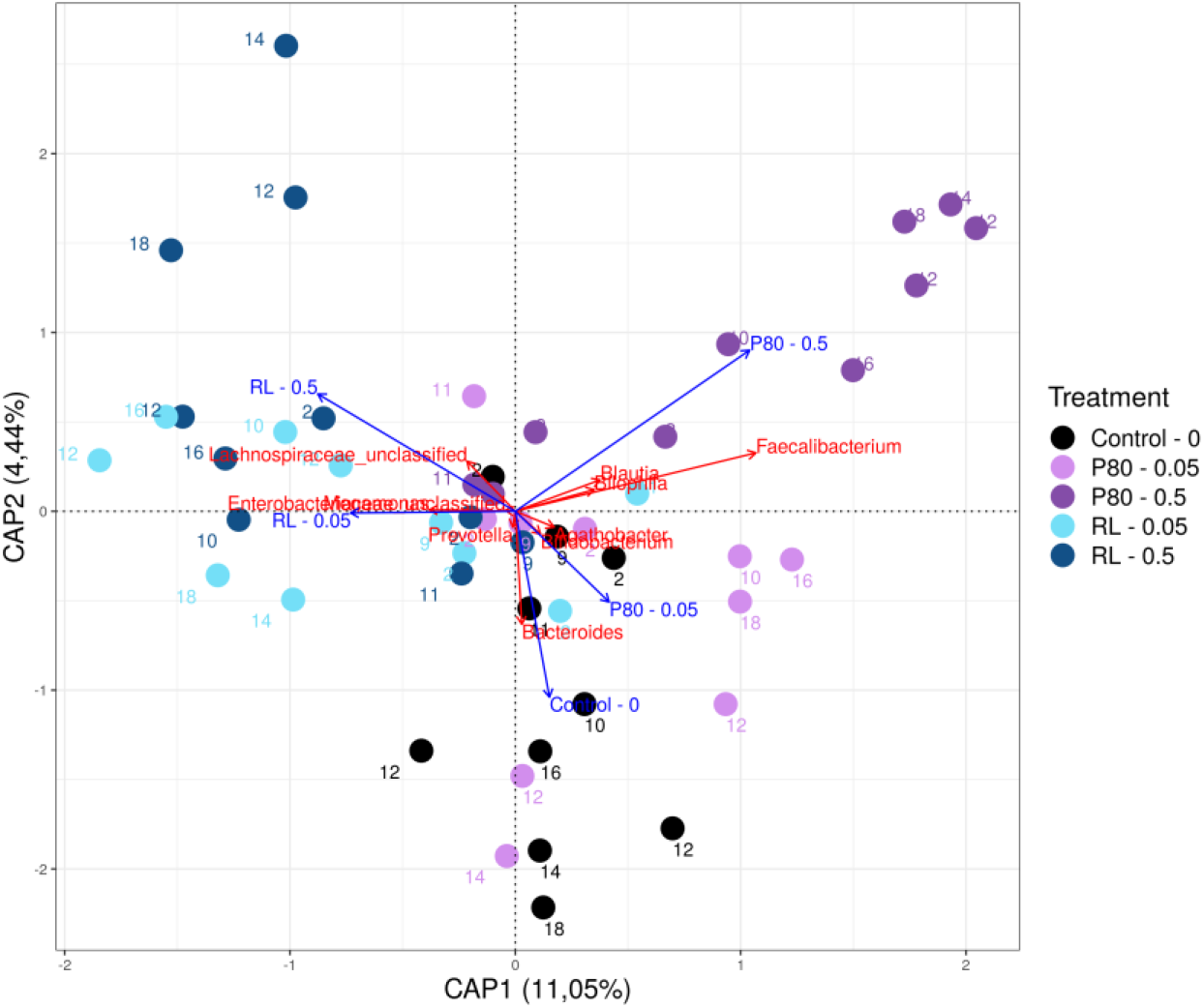
Type II scaling triplot obtained from partial distance based redundancy analysis for the absolute abundances of microbial genera in luminal suspensions from a 16 and 18 day SHIME experiment investigating the impact of a 7 day treatment with TWEEN80 and rhamnolipids (0,05 m% and 0,5 m%) on the gut microbiota from two human faecal donors. Absolute abundances were retrieved by use of 16S rRNA amplicon sequencing and flow cytometry. The different treatment conditions were set as explanatory variables (blue arrows) and the 5 most prevalent genera were set as response variables (red arrows). Factors Donor and Timepoint were partialled out.

Calculation of the Shannon, Simpson and Chao1 indices indicated that neither TWEEN80 nor rhamnolipids showed remarkable or consistent impacts on the richness or diversity of the microbial communities. Only the Chao1 index was lowered with the rhamnolipid treatments in both lumen and mucus (Supplementary Figures 4-9).

Next, Deseq2-analysis was employed to explore which other bacterial genera were altered by the addition of TWEEN80 or rhamnolipids. TWEEN80 was shown to increase the relative abundance of 14 genera and lower the abundance of 10 genera in the luminal environment. TWEEN80 also inflated 18 genera in the mucosal environment as well as suppressing 11 others. Rhamnolipids on the other hand stimulated 10 and suppressed 48 genera in the lumen and they increased 6 genera and lowered 56 genera in the mucus.

Of the genera that were increased by both TWEEN80-concentrations in the luminal environment for both donors, *Erysipelatoclostridium* (LFC ≥ 1,74; Padj ≤ 0,019), *Collinsella* (L2FC ≥ 1,59; Padj ≤ 0,051), *Hungatella* (L2FC ≥ 1,72; Padj ≤ 0,019) were the three that presented with the strongest increase in abundance, while *Alistipes* (L2FC ≤ −1,52; P_adj_ ≤ 0,040) was the genus that was the most suppressed (Figure 5;Supplementary Figure 11; Supplementary Table 1 & 2). For the rhamnolipids, the most stimulated genus in the luminal suspension was unclassified Enterobacteriaceae (L2FC ≥ 2,3; Padj ≤ 0,046), and the three most suppressed genera were *Faecalibacterium* (L2FC ≤ −12,27; P_adj_ ≤ 0,001), *Blautia* (L2FC ≤ −5,46; P_adj_ ≤ 0,001) and unclassified Ruminococcaceae (L2FC ≤ −5,95; P_adj_ ≤ 0,001) (Figure 5;Supplementary Figure 11; Supplementary Table 1 & 2).

**Figure 5:**
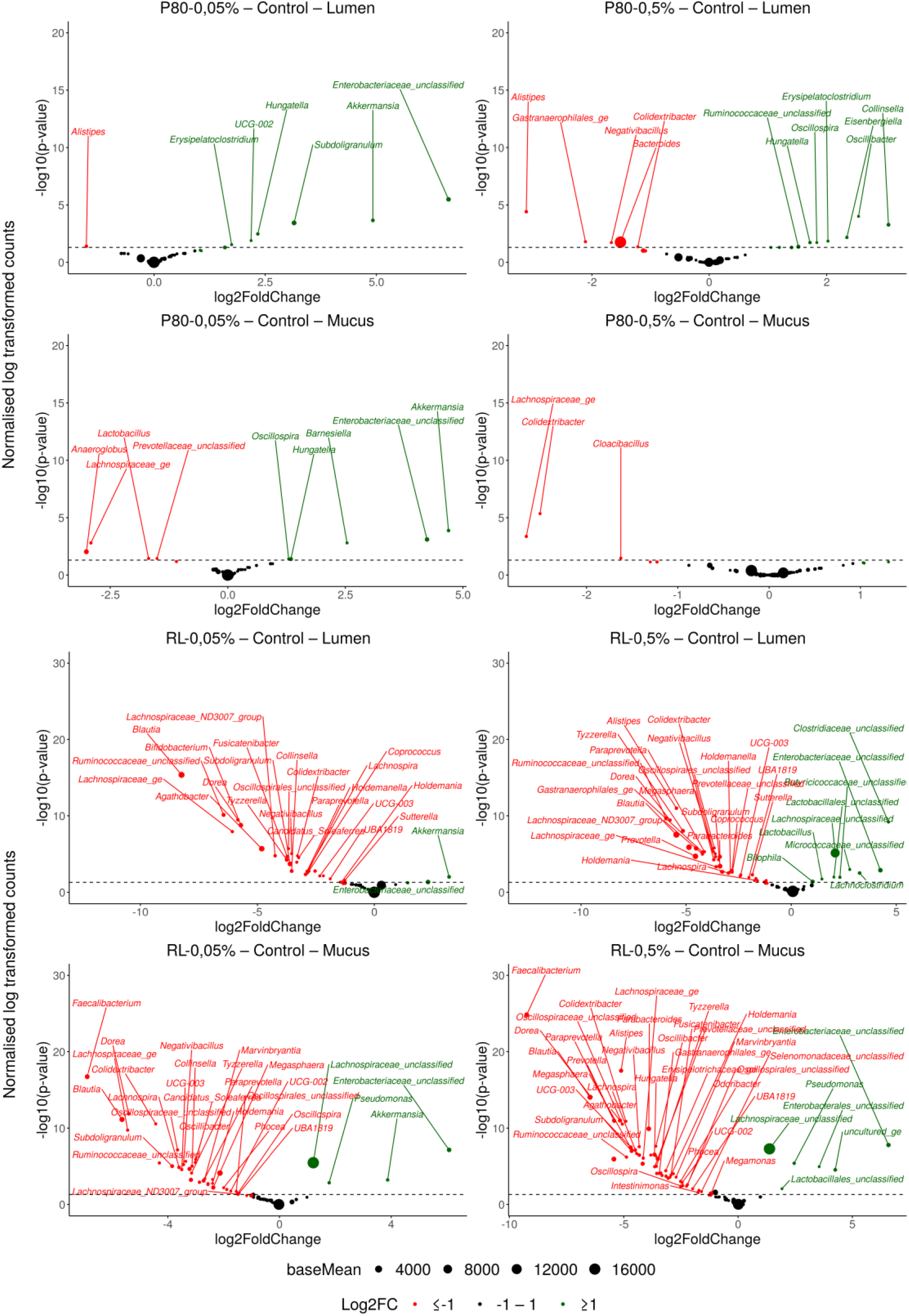
Volcano plots designating shifts to the luminal and mucosal microbial communities in response to treatment with TWEEN80 and rhamnolipids (0,05 m% and 0,5 m%) during a 16 and an 18 day SHIME experiment investigating the impact of a 7 day treatment with rhamnolipids on the gut microbiota from two human faecal donors. Log2FoldChange (L2FC) of genus abundances for the treatments versus the controls are presented on the x-axis and the log-transformed adjusted p-value is indicated on the y-axis. Significantly in- or decreased genera are indicated in green and red respectively. The dashed line represents the significance threshold of α = 0,05.

In the mucosal compartment, both concentrations of TWEEN80 significantly suppressed the genus of Lachnospiraceae_ge (L2FC ≤ −2.66; P_adj_ ≤ 0,001). The top three genera increased in abundance by both concentrations of rhamnolipids were the unclassified Enterobacteriaceae (L2FC ≥ 6,07; Padj ≤ 0,001), the unclassified Lachnospiraceae (L2FC ≥ 1,21; Padj ≤ 0,001) and *Pseudomonas* (L2FC ≥ 1,78; Padj ≤ 0,001) (Figure 5; Supplementary Figure 12; Supplementary Table 1&2). The top three most suppressed genera by rhamnolipids in the mucosal compartment were *Faecalibacterium* (L2FC ≤ −6,89; P_adj_ ≤ 0,001), *Blautia* (L2FC ≤ −5,64; P_adj_ ≤ 0,001) and *Dorea* (L2FC ≤ −5,4; P_adj_ ≤ 0,001) (Figure 5; Supplementary Figures 12; Supplementary Table 1&2).

### 3.4 Short Chain Fatty Acids

Short chain fatty acid levels were measured throughout the experiments in the luminal compartments of the SHIME to assess the general activity of the gut microbiota in response to the emulsifiers. In these experiments, TWEEN80 and rhamnolipids had only limited effects on the total SCFA concentrations (Supplementary Figure 13). However, individual SCFA-levels were still shifted and this in an opposite way for each emulsifier. Propionate levels were lowered by the addition of TWEEN80, especially for donor 2, but they were increased in the presence of rhamnolipids (Figure 6, Table 1). On the other hand, butyrate levels were increased by TWEEN80, but were decreased by rhamnolipids compared to the end of the stabilisation phase (Figure 6, Table 1). Acetate levels weren’t seemingly affected by either emulsifier, but since the cell counts were reduced by an order of magnitude by the rhamnolipids, acetate production must have increased (Supplementary Figure 14). Lastly, the high and low responder status for which the faecal donors were selected were clearly visible here.

**Figure 6:**
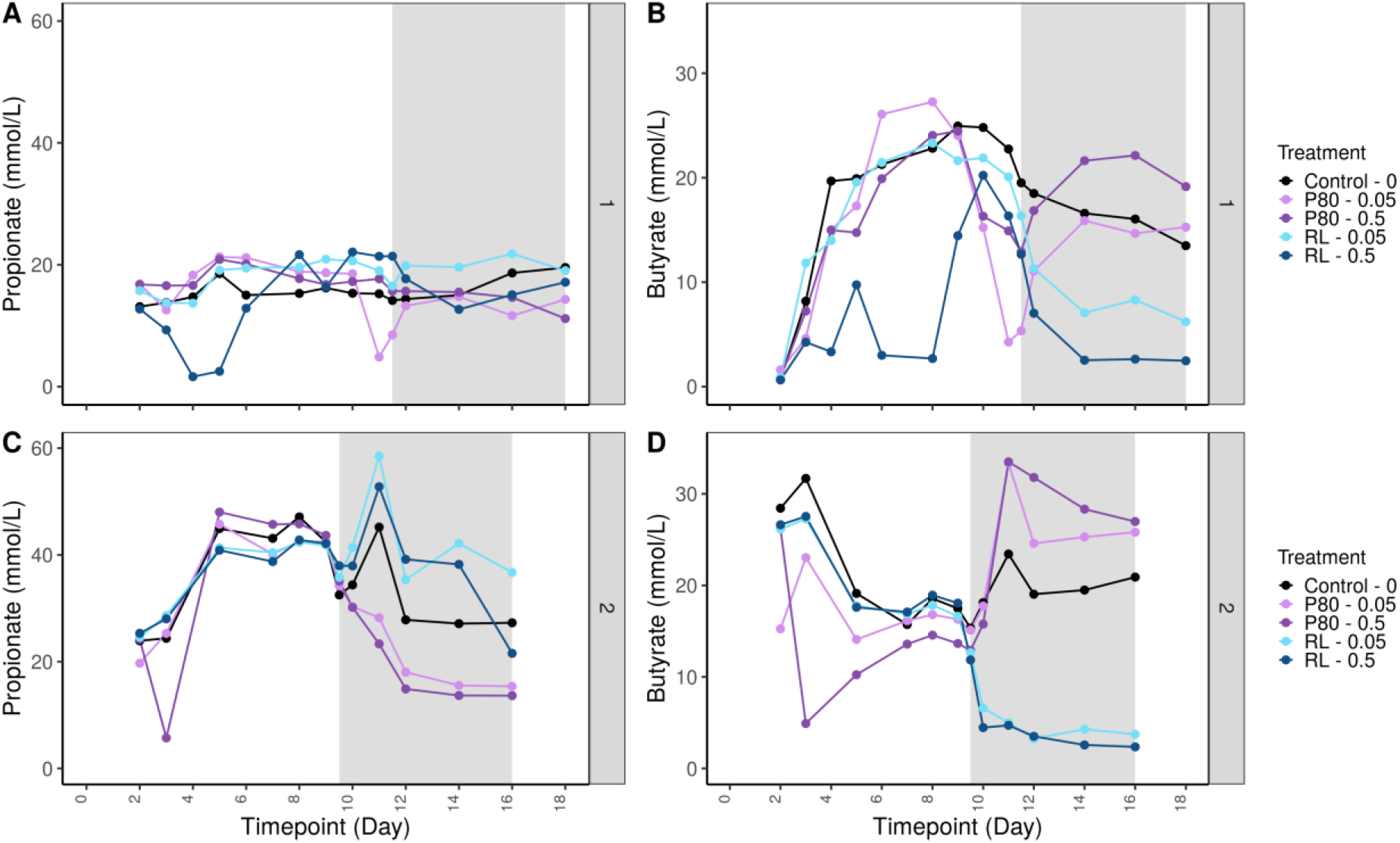
Propionate (A and C) and butyrate levels (B and D) (mM) in luminal suspension during a 16 and 18 day SHIME experiment investigating the impact of TWEEN80 and rhamnolipids (0,05 m% and 0,5 m%) on the gut microbiota of two human faecal donors. Donor 1 was selected for low emulsifier sensitivity and donor 2 for high emulsifier sensitivity. The 7-day treatment period is indicated by the grey background.

Given the obvious alterations in the absolute abundance of *Faecalibacterium,* the correlation of its abundance with the levels of butyrate were checked. The butyrate levels correlated significantly with the abundance of *Faecalibacterium* for both TWEEN80 (Corr_0,05% = 0,51; p_0,05% = 0,02; Corr_0,5% = 0,64; p_0,5% = 0,003) and the rhamnolipids (Corr_0,05% = 0,77; p_0,05% = 7,63e-5; Corr_0,5% = 0,74; p_0,5% = 0,0002).

### 3.5 Metabolomics

Both PCA on the untargeted dataset, as well as PCoA on the targeted dataset showed that rhamnolipids caused a clear shift in the metabolome of both donors but that TWEEN80 didn’t cause very prominent changes (Figure 7). Nevertheless, statistical analysis using the MetaboAnalyst website unveiled several significantly altered metabolites for both emulsifiers. For TWEEN80, both concentrations significantly decreased the levels of 3-indoleacetic acid and 2-aminobutyric acid and significantly increased the levels of methylcyclohexanecarboxylate (Figure 8). For rhamnolipids, hypoxanthine levels were significantly decreased and levels of N-acetyl-L-methionine, N-acetylglucosamine, beta-pinene, ethylbenzene and L-arginine were significantly increased by both concentrations and in both donors (Figure 9).

**Figure 7:**
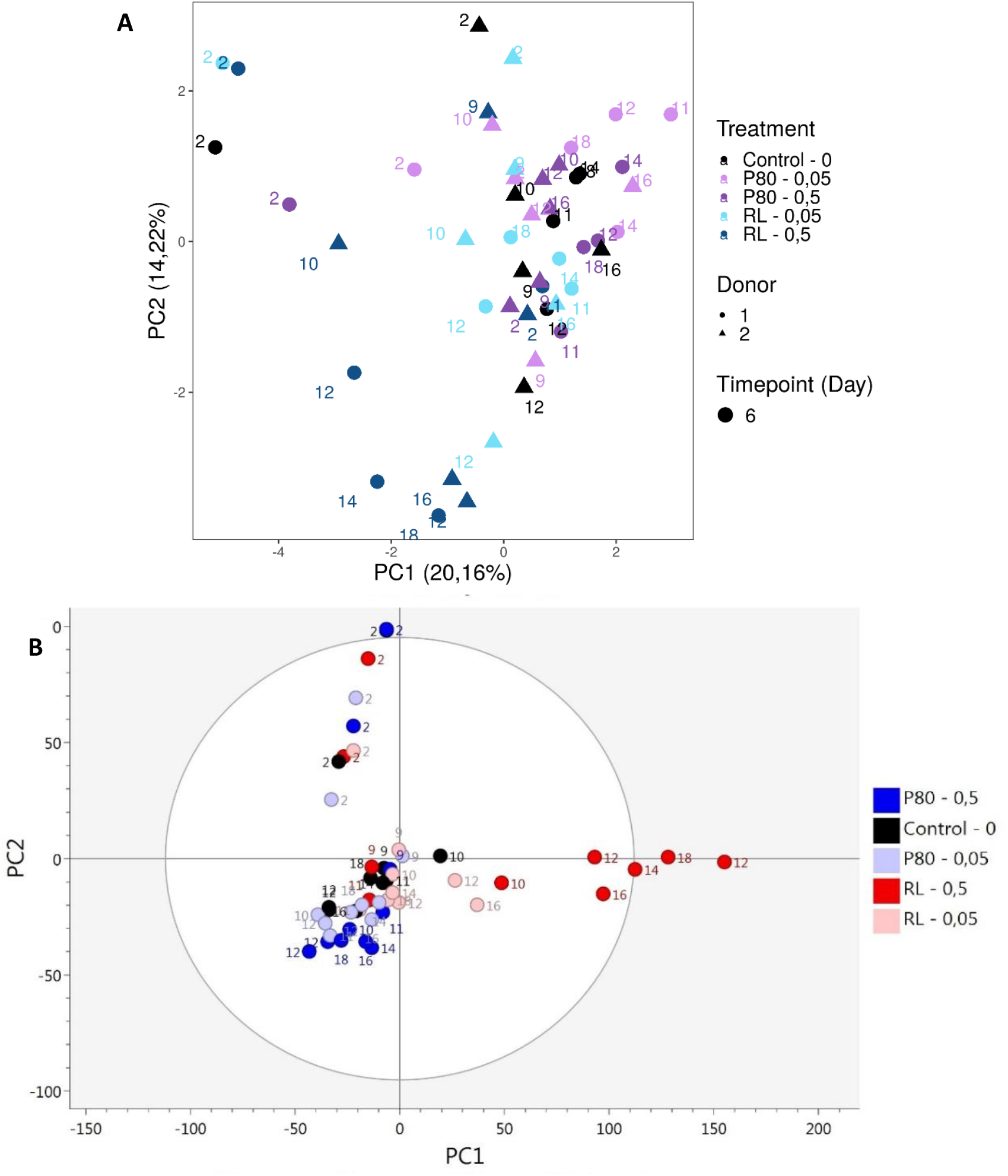
Principle coordinate analysis of targeted (A) and principle component analysis of untargeted (B) metabolomics data extracted from luminal suspensions from a 16 and an 18 day SHIME experiment investigating the impact of TWEEN80 and rhamnolipids (0,05 m% and 0,5 m%) on the gut microbiota from two human faecal donors.

**Figure 8:**
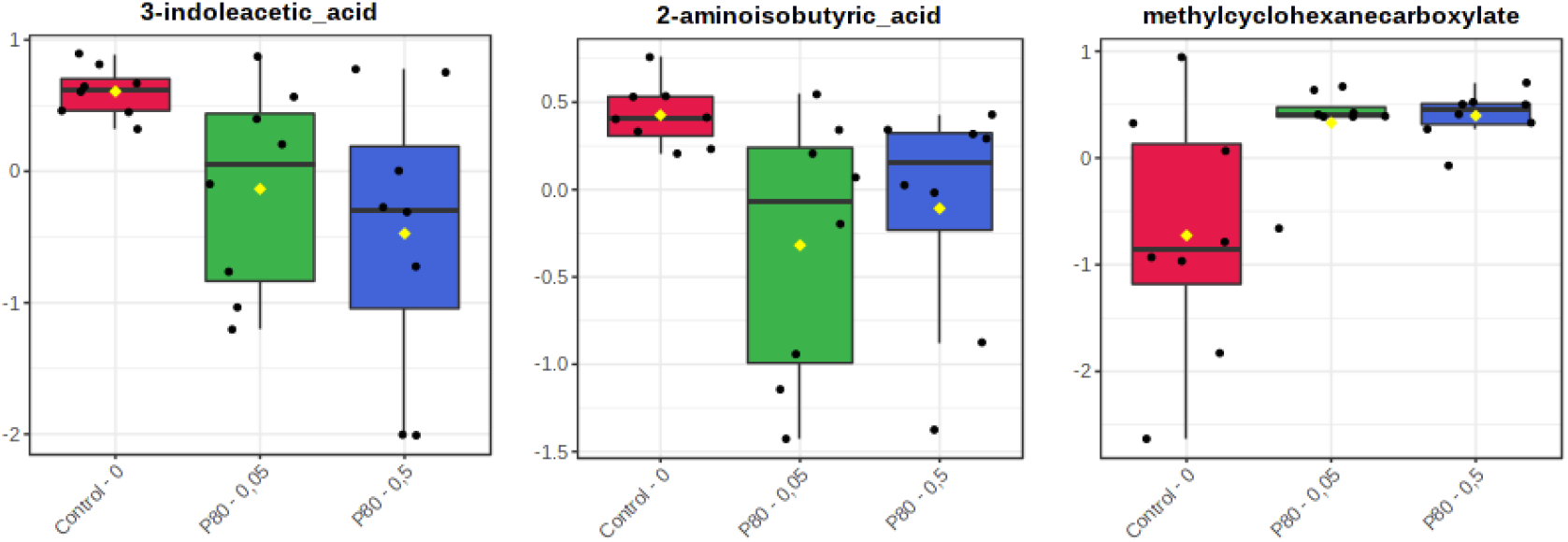
Microbial metabolites from the targeted metabolite dataset that were significantly affected by both treatments with TWEEN80 (0,05% and 0,5%) in luminal suspensions from a 16 and an 18 day SHIME experiment investigating the impact of TWEEN80 on the composition and functionality of the gut microbiota from two human faecal donors. Compounds were considered relevant when they were indicated as significantly different by the Wilcoxon Rank-Sum test, the VIP value was larger than 1 in the PLS-DA and if p(1) >0,1 and p(corr)[1] > 0,4 in the OPLS-DA for treatment with both TWEEN80 concentrations.

**Figuur 9:**
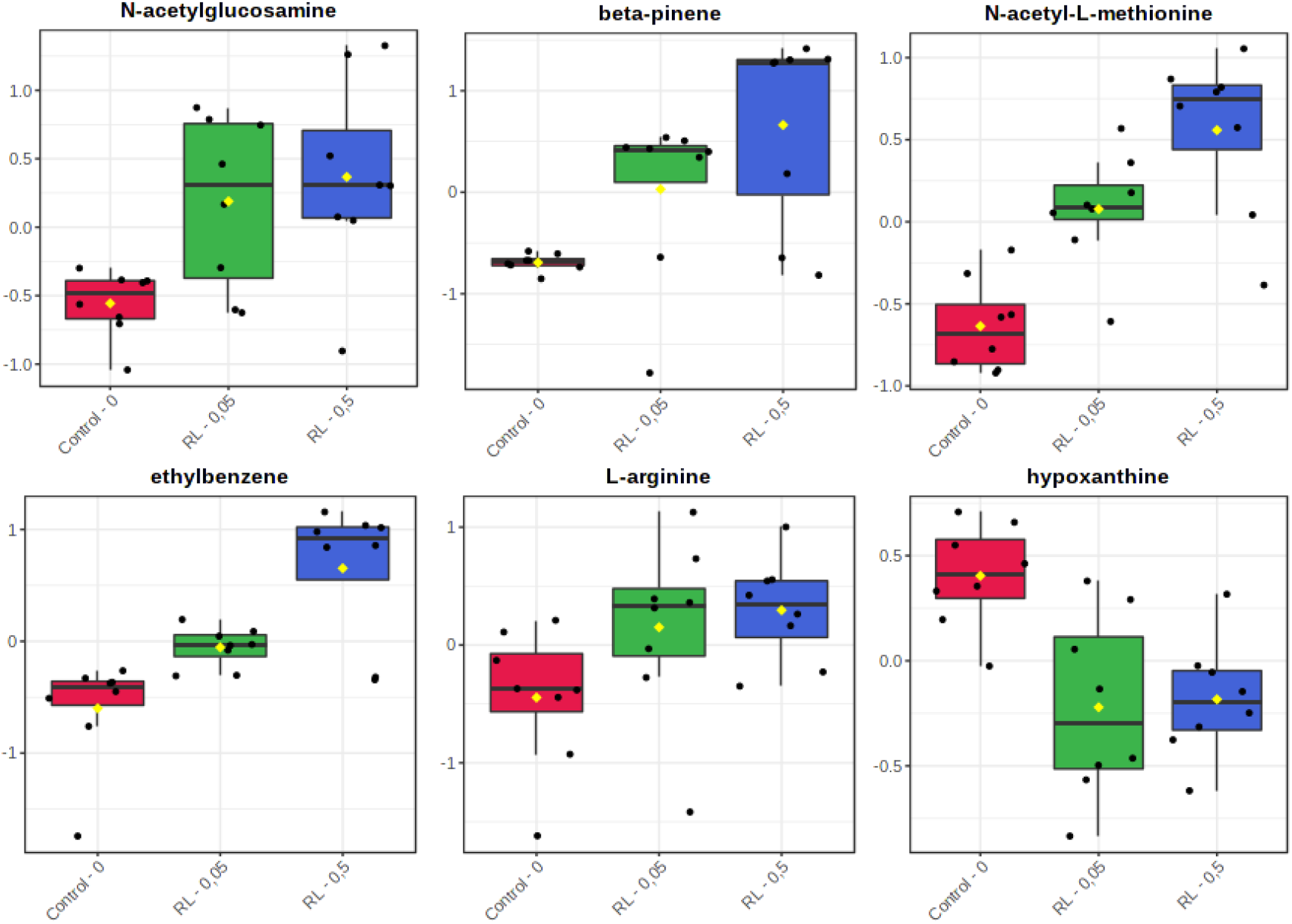
Microbial metabolites from the targeted metabolite dataset that were significantly affected by both treatments with rhamnolipids (0,05% and 0,5%) in luminal suspensions from a 16 and an 18 day SHIME experiment investigating the impact of rhamnolipids on the composition and functionality of the gut microbiota from two human faecal donors. Compounds were considered relevant when they were indicated as significantly different by the Wilcoxon Rank-Sum test, the VIP value was larger than 1 in the PLS-DA and if p(1) >0,1 and p(corr)[1] > 0,4 in the OPLS-DA for treatment with both rhamnolipid concentrations.

Furthermore, the enrichment analysis tool on the MetaboAnalyst website indicated that TWEEN80 caused shifts in the metabolome which pointed towards a state of obesity and arthritis (Supplementary Figures 15, 19). These shifts were significant (p < 0,05 for obesity and p < 0,02 for arthritis) for both TWEEN80-concentrations and were based on the metabolites pictured in the Supplementary Figures 16-18 and 20-21. The metabolomic shifts caused by the rhamnolipids were only significant for the higher concentration, but also these were reminiscent of obesity (p < 0,07), as well as arthritis (p < 0,1) and irritable bowel syndrome (p<0,009)(Supplementary Figures 22-30).

For the untargeted metabolomics dataset, significantly altered metabolites were also sought using both the S-plots (Supplementary Figure 32, 33) and VIP-scores in SIMCA as well as statistical analysis in MetaboAnalyst. MetaboAnalyst detected a vast set of compounds. SIMCA did not identify any meaningful compounds.

## 4. Discussion

In this paper, two human faecal microbiota that were previously selected based on their susceptibility to dietary emulsifiers, were exposed for one week to TWEEN80 and rhamnolipids in the M-SHIME system. The results partially confirmed the observations from our previous 48h batch incubations. While during the 48h batch incubations, TWEEN80 caused no significant trends in microbial composition or functionality, in the SHIME-system, both TWEEN80 and rhamnolipids altered the composition and functionality of the gut microbiota from the two faecal donors. The overall impact was greater for the rhamnolipids than for TWEEN80, and was dependent on the composition of the original microbial community, as well as the intestinal micro-environment. The effects also reached an equilibrium after about 3 days of exposure.

TWEEN80 and rhamnolipids caused both opposing and similar effects in the gut microbiota. As such, TWEEN80 increased cell counts, lowered propionate levels, increased butyrate levels and increased the abundance of several genera of bacteria: *Faecalibacterium, Agathobacter, Subdoligranulum, Blautia, Anaerostipes, Oscillobacter, Oscillospira* and *unclassified Ruminococcaceae*. Rhamnolipids caused the opposite efects. Why certain bacterial genera experience an advantage from the presence of TWEEN80 is unclear at this point. This advantage might have stemmed from TWEEN80 providing a source of nutrition, it altering the availability/consistency of the SHIME-feed or the emulsifier might have altered the crosstalk between the different species. Similarities in the effects of the emulsifiers were that both compounds increased the relative abundance of the genera *Akkermansia muciniphila* (only for the 0,05 m% conditions) as well as unclassified Enterobacteriaceae. Both also altered the level of microbial metabolites that were previously related to obesity, arthritis and instestinal issues.

Rhamnolipids have not previously been investigated with respect to the gut microbiota, but a number of other studies have researched the impact of TWEEN80 on the gut microbiota before. Their findings contradict ours to a certain degree. Firstly, Naimi et al., (2021) have found that TWEEN80 decreased the relative abundance of *Faecalibacterium, Akkermansia*, unclassified Ruminococcaceae and *Oscillospira*, whereas we’ve found those genera to be increased. They also detected an increased abundance of Enterobacteriaceae, as well as a slight increase in bacterial density, as measured through 16S rRNA qPCR, which does agree with our findings. Secondly, Chassaing et al 2017 have found, also in a SHIME-setting, that TWEEN80 did not alter the levels of SCFA significantly (Chassaing 2017), whereas we’ve found alterations in propionate and butyrate levels. Lastly, Singh & Ishikawa, (2016) have found that gavaging C57BL/6 mice with 1% of of their bodyweight in TWEEN80 reduced the levels of all three SCFA in the murine feces, decreased bacteroidetes abundance (which we also observed) and increased the abundance of opportunistic pathogens. In short, the effects of TWEEN80 on the gut microbiota show little consistency in literature. This may be due to the use of different models and set-ups or differences in the microbial community at the start of the experiments.

During the SHIME experiments, the low or high emulsifier sensitivity, for which the microbiota were selected, was reproduced. This was particularly visible in the SCFA data, for which donor 2 showed more pronounced alterations than donor 1 during both the 48h batch incubations and the SHIME experiments. In the cell count, amplicon sequencing and metabolomics data the responder status was less visible. Nevertheless, the PCA of the untargeted metabolomics data, showed that the response of the microbiota from donor 2, the high responder, was greater than that of donor 1 (Supplementary Figure 31).

Some of our results can be discussed in relation to host health. A decrease in cell concentrations, for instance, which we observed after exposure to rhamnolipids, has recently been related to Crohn’s disease [41]. The effects of the emulsifiers on propionate and butyrate may also have consequences for a host. Both of these SCFA are considered health-promoting, but their impact on physiology is clearly different. Butyrate is mainly of importance as an energy source and a signaling molecule for the gut epithelium, while propionate reaches the liver and periferal tissues and there helps regulate blood glucose and cholesterol metabolism [10]. Since each of the emulsifiers only caused the ratio’s of butyrate and propionate to switch, rather than to decrease only one of them, it is hard to assess what the effect on host health might be, should this pattern repeat itself *in vivo*.

The potential impact of the shifts in microbial composition isn’t straighforward either. The butyrate levels could for each emulsifier be correlated to the abundance of *Faecalibacterium*, which is considered beneficial for its butyrate producing capacities cancer [42–44]. *Akkermansia muciniphila* is also considered health-promoting, mainly because of its interaction with the hosts mucus layer [45,46], so its increased abundance with the lower concentrations of emulsifiers are, in essence, positive. At the higher emulsifier concentrations, the logfoldchange of *Akkermansia* was much lower though (Supplementary tables 1&2), so the effect was not concentration dependent. The increase in unclassified Enterobacteriaceae after the addition of both TWEEN80 and rhamnolipids is generally considered disadvantageous [47–49]. In short, the emulsifiers caused both health promoting and health-adverse microbial alterations.

Little can be said about the health impact of the trends in the metabolome that were caused by the emulsifiers, because the metabolic role of many of the detected metabolites is not known. MetaboAnalyst detected potential relationships of metabolite shifts with disease patterns, but for many of metabolites these conclusions were based on, the mechanistic relationship with the disease is not clear. The decrease in vaccenic acid we observed, for instance, cannot be reliably linked to obesity, because the SHIME-model did not simulate potential absorption of this compound, while the studies that linked vaccenic acid to obesity were performed *in vivo* [50,51]. Indole acetic acid is an exception though. This compound was lowered in the presence of the emulsifiers and this trend was related by MetaboAnalyst to enthesitis-related arthritis, a type of Spondyloarthritis. For this metabolite, the molecular pathways are known. It can be generated through microbial tryptophane (Trp) metabolism and is known to provide potent anti-inflammatory effects, as well as regulate the hosts immune system [52]. In arthritis patients, faecal Trp metabolites and microbial Trp metabolism pathways were indeed reduced [52]. The lowering of indole acetic acid as a consequence of the emulsifier treatments could thus be considered disadvantageous to the host.

Rhamnolipids were included in this research to assess whether they could provide a safer, clean-label alternative to the conventional emulsifier TWEEN80 with regards to intestinal health. When comparing the current results for rhamnolipids and TWEEN80 in this context it must be stressed that it’s effects on the gut microbiota were more severe at similar concentrations. Rhamnolipids lowered butyrate levels, increased levels of potentially pathogenic Enterobacteriaceae and reduced the abundance of many other microbial genera, thereby reducing the microbial diversity. Microbial diversity in the intestine is a factor that is consistently linked with positive health outcomes [41]. In that sense, utilizing rhamnolipids as novel dietary emulsifiers must be approached with caution. It is possible though, that a lower concentration of rhamnolipids can be used in food products, because of better emulsifying capacities, and that the effects thus become less severe.

A note that needs to be made is that there is still no certainty about whether the effects we observed here for TWEEN80 and rhamnolipids might also occur in a human colon. It is clear from our studies, as well as the literature, that the model used to investigate the impact of dietary emulsifiers strongly dictates the outcomes. Up till now, *in vivo* research regarding emulsifiers mainly focused on host health endpoints. New *in vivo* research should also focus on the metagenomic and metabolomic impact of the gut microbiota. On top of model-related differences, the interindividual variability between the microbiota from different donors also affects the potential impact of the emulsifiers. The differences in response from people of different sexes, ages, ethnicities, living area’s, etc. can thus still be characterized by including a larger variety of human faecal donors into experiments like these.

There are several other points that also deserve further study. The resilience of the gut microbiota (i.e. its ability to return to a previous state after a disturbance), for instance, was not tested in our SHIME experiments. This can easily be investigated by including a wash-out period at the end of the current set-up. Next, it would be useful to investigate which bacteria are responsible for specific alterations in the metabolome. Lots of microbial metabolic pathways are already known, but in an interacting consortium of bacteria, there is still no overview over who does what. Lastly, most of the alterations observed in this study cannot yet be linked to effects in the host, let alone health outcomes. Even though the metabolomics analyses indicated metabolomic shifts in the direction of obesity, arthritis and gut inflammatory diseases, the precise mechanisms behind most of these indications is still missing.

In summary, the current paper describes the opposing effects of TWEEN80 and rhamnolipids on the gut microbiota from two faecal donors, previously selected based on their response status. Both emulsifiers altered the composition and functionality of the gut microbiota, with the effects of the rhamnolipids being more severe and potentially undesirable in a gut health context. In this sense, the rhamnolipids may not necessarily form a safer clean label alternative for TWEEN80. The individuality of the microbiota, for which they were selected, was largely preserved during the SHIME experiments. Further research should focus on assessing *in vivo* impacts of emulsifier exposed gut microbiota, establishing the mechanisms behind the species-specific effects and identifying whether the alterations to the gut microbiota are final or reversible.

## 6. Supplementary materials and methods

**Figure 1:**
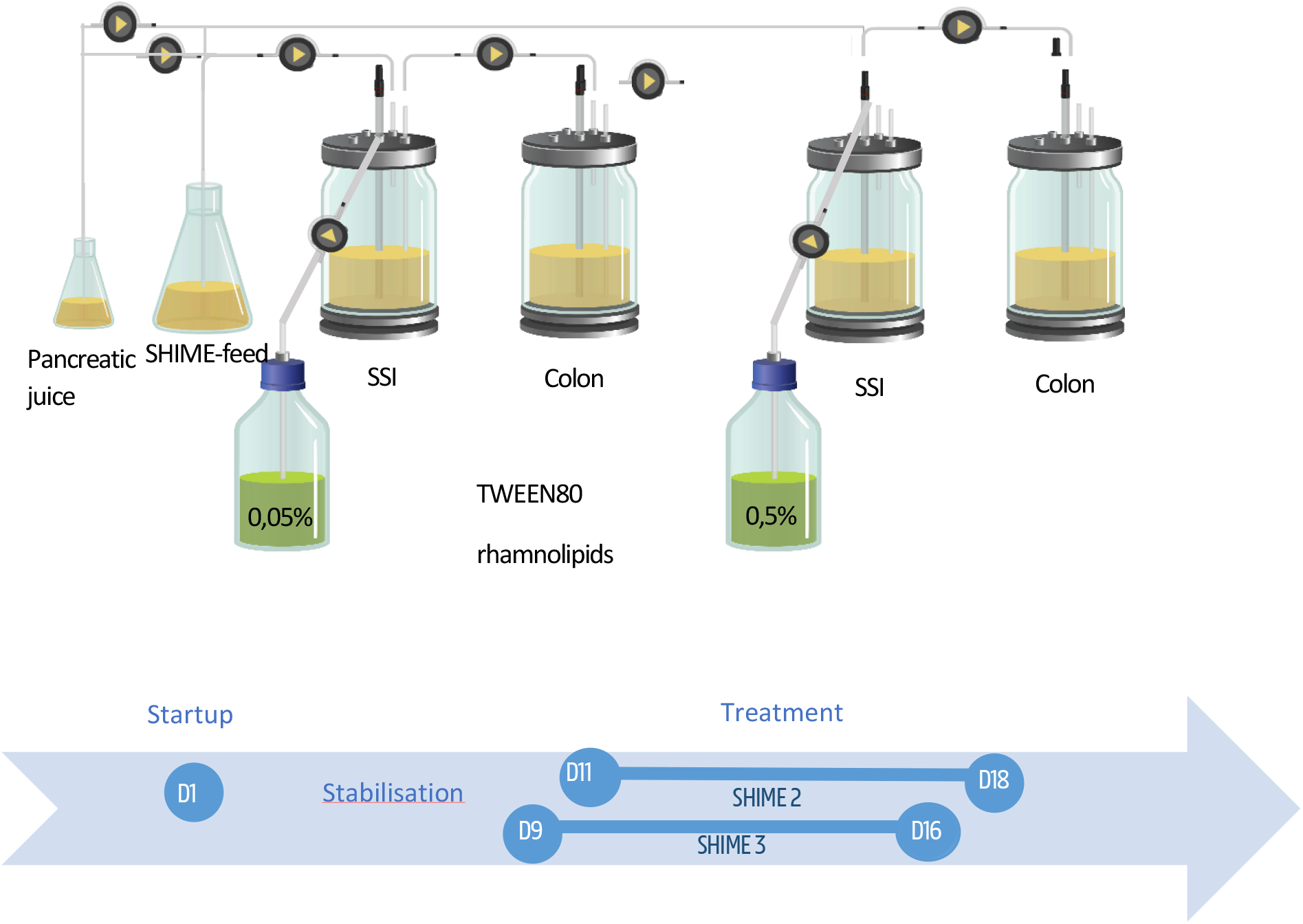
Schematic representation of an experiment with the Simulator of the Human Intestinal Microbial Ecosystem (SHIME) investigating the impact of 0,05 m% and 0,5 % TWEEN80 and rhamnolipids on the composition and functionality of the gut microbiota from 2 human faecal donors.

**Table 1:** Chemical composition of nutritional SHIME-feed.

| Component | Concentration (g/L) |
| --- | --- |
| arabinogalactan | 1,2 g/L |
| Pectin | 2 |
| Xylan | 0,5 |
| Glucose | 0,4 |
| Yeast extract | 3 |
| Special Pepton | 1 |
| Mucin | 2 |
| Cystein-HCl | 0,5 |
| Starch | 4 |

**Table 2:** Time table of pumping action between different SHIME-compartments.

| Feed =><br>Stomach/small<br>intestine | Pancreatic juice =><br>Stomach/small<br>intestine | Emulsifier =><br>Stomach/small<br>intestine | Stomach/small<br>intestine => Colon | Colon => Waste |
| --- | --- | --- | --- | --- |
| 9u -> 9u30 | 10u -> 9u15 | 9u50 -> 10u | 12u30 -> 14u30 | 13u30 -> 15u30 |
| 17u -> 17u30 | 18u -> 17u15 | 18u50 -> 19u | 20u30 -> 22u30 | 21u30 -> 23u30 |
| 1 -> 1u30 | 2 -> 2u15 | 2u50 -> 3u | 4u30 -> 6u30 | 5u30 -> 7u30 |

**Table 3:**
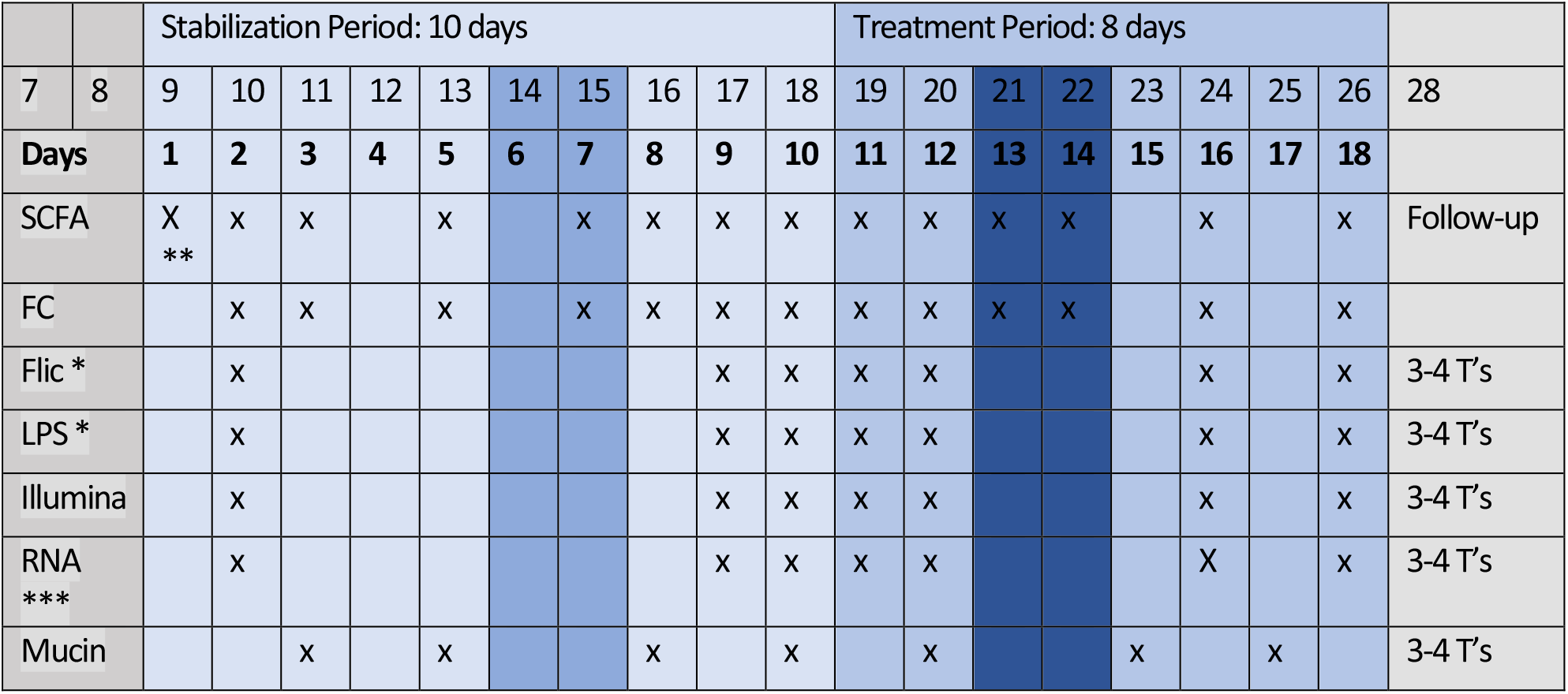
Sampling scheme for SHIME 1 with donor 1.

**Table 4:**
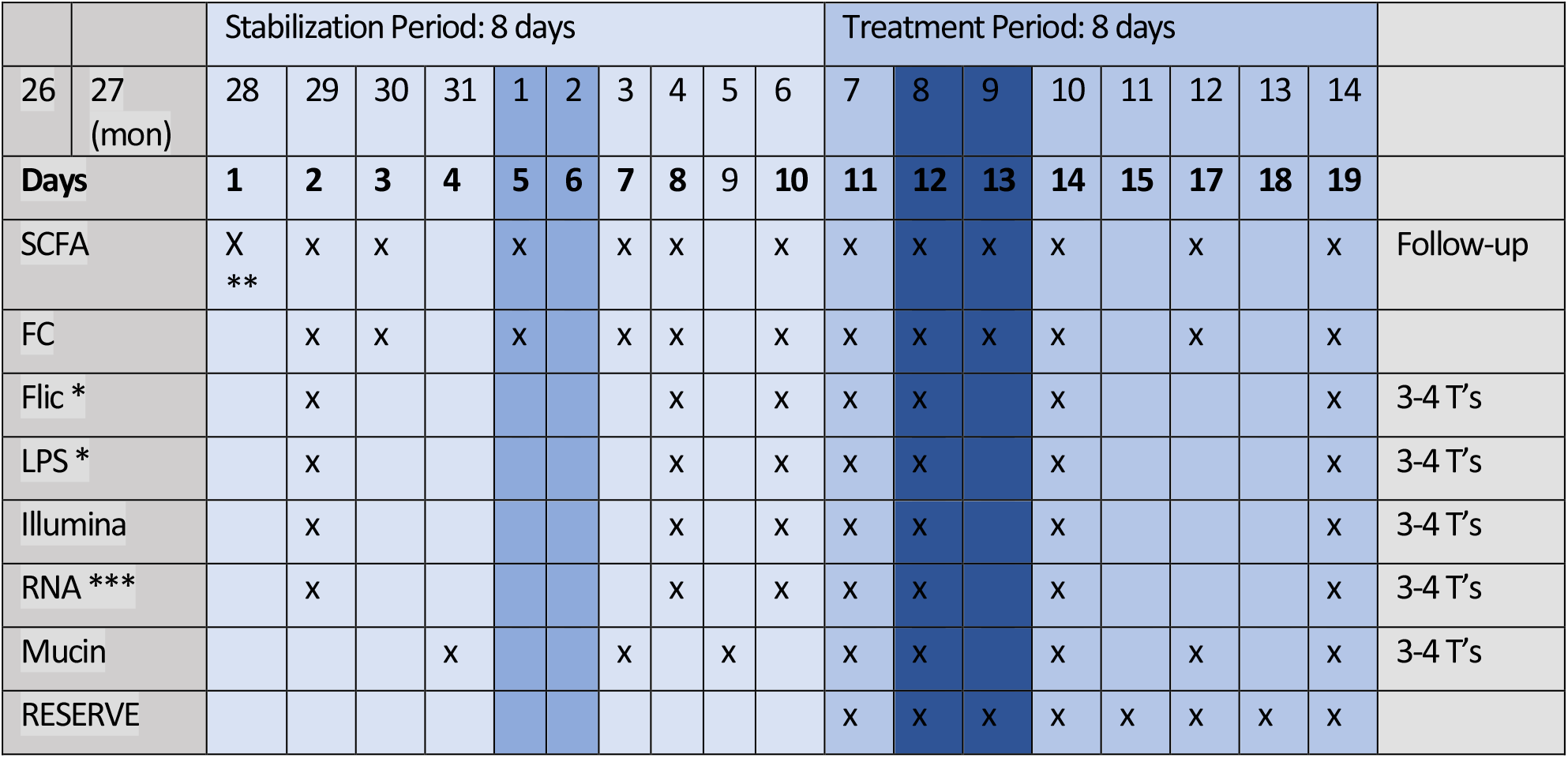
Sampling scheme for SHIME 2 with donor 2.

