## Supplementary Figures for "Differential Impact of rhamnolipids and TWEEN80 on composition and functionality of the gut microbiota in the SHIME system"

#### 1. Cell counts

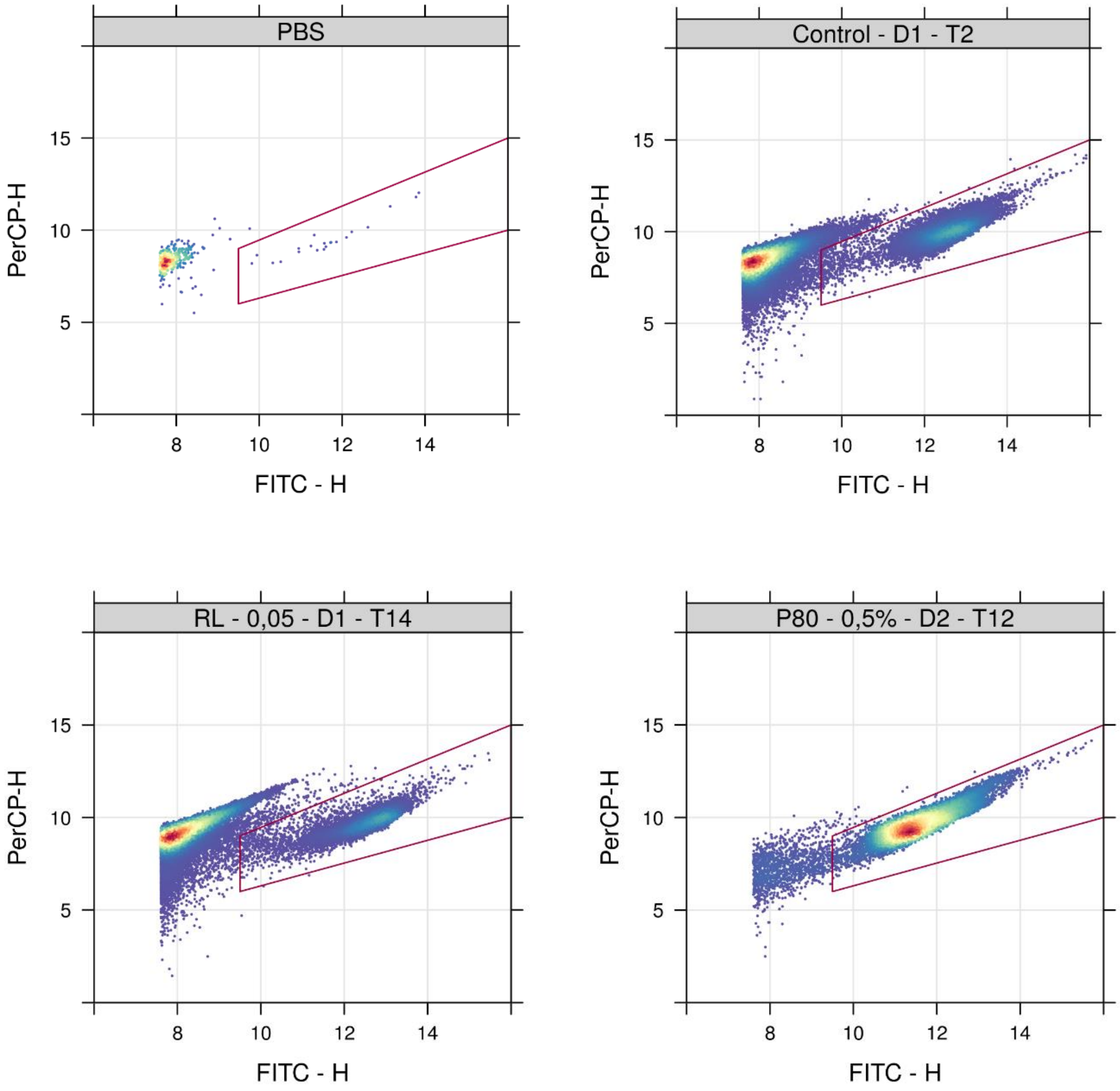


*Figure 1: Density plots of cell counts obtained through SYBRgreen staining and flow cytometry of several random samples from luminal suspensions from a 16 and 18 day SHIME experiment investigating the impact of TWEEN80 and rhamnolipids (0,05 m% and 0,5 m%) on the gut microbiota of two human faecal donors with a 7 day treatment period. The red lines indicate the gating utilized to obtain total cell concentrations in R (version 4.1.0).*

#### 2. Amplicon sequencing

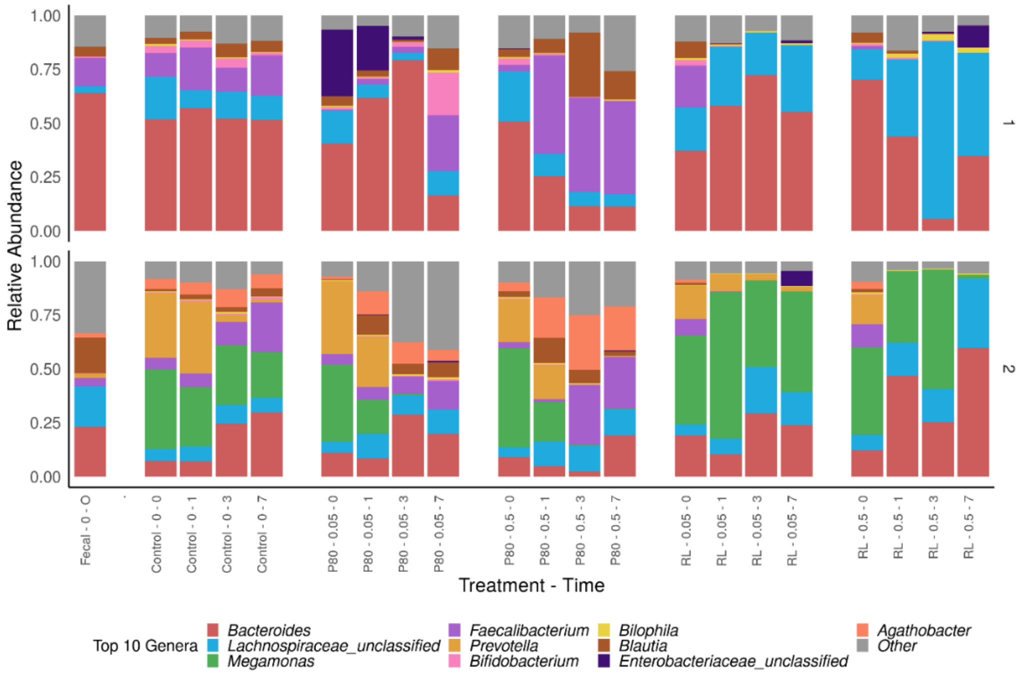


*Figure 2: Relative abundances of the 10 most abundant genera derived from 16S rRNA gene amplicon sequencing, measured in luminal suspensions of a 16 and 18 day SHIME experiment investigating the impact of TWEEN80 and rhamnolipids on the gut microbiota from two human faecal donors. For each condition, different timepoints are indicated relative to the start of the treatment: timepoint 0 (days).*

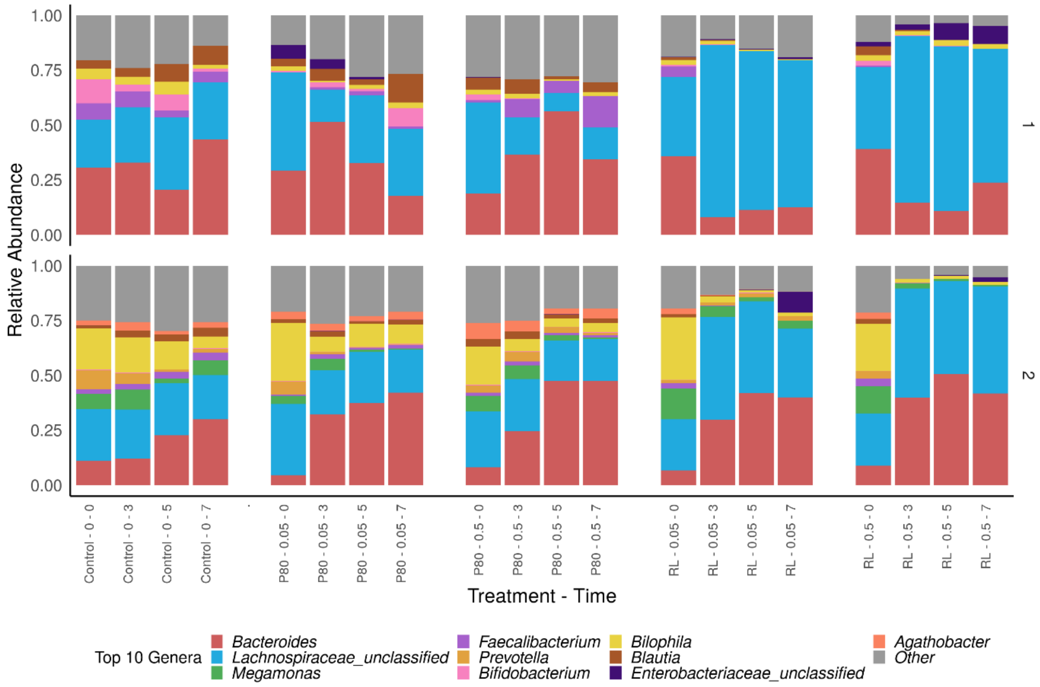


*Figure 3: Relative abundances of the 10 most abundant genera derived from 16S rRNA gene amplicon sequencing, measured in the mucosal compartments of a 16 and 18 day SHIME experiment investigating the impact of TWEEN80 and rhamnolipids on the gut microbiota from two human faecal donors. For each condition, different timepoints are indicated relative to the start of the treatment: timepoint 0 (days).*

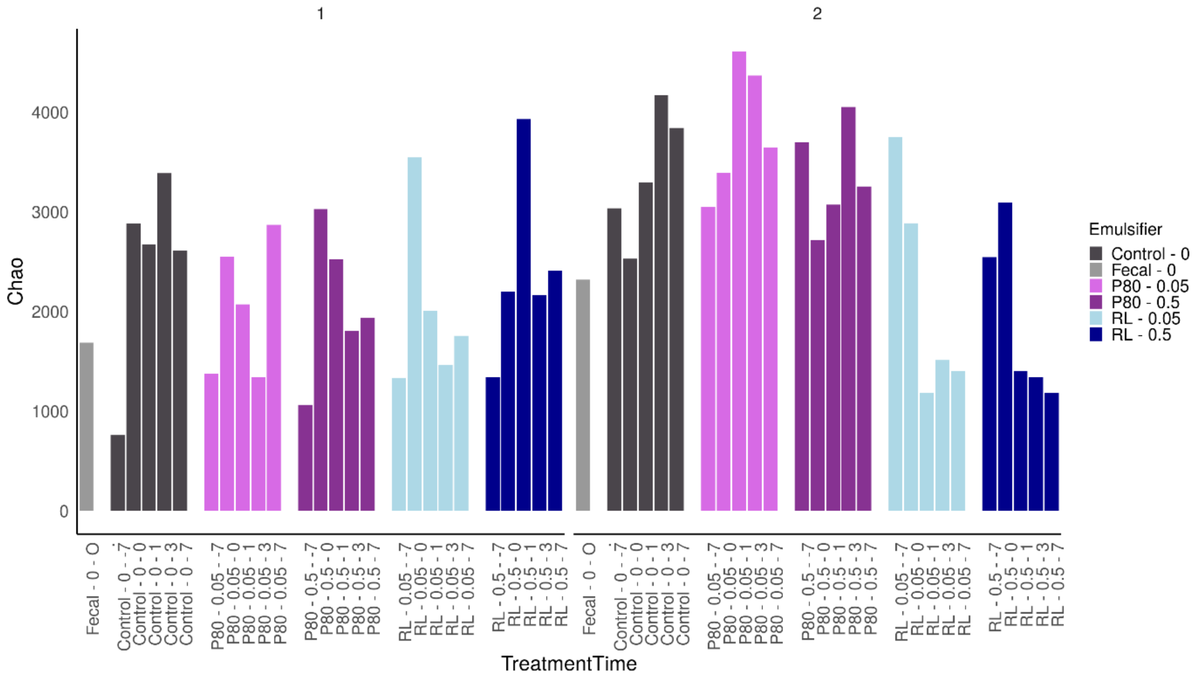


*Figure 4: Chao richness of the simulated gut microbiota in luminal suspensions from a 16 and 18 day SHIME experiment investigating the impact of TWEEN80 and rhamnolipids (0,05 m% and 0,5 m%) on the gut microbiota from two human faecal donors.*

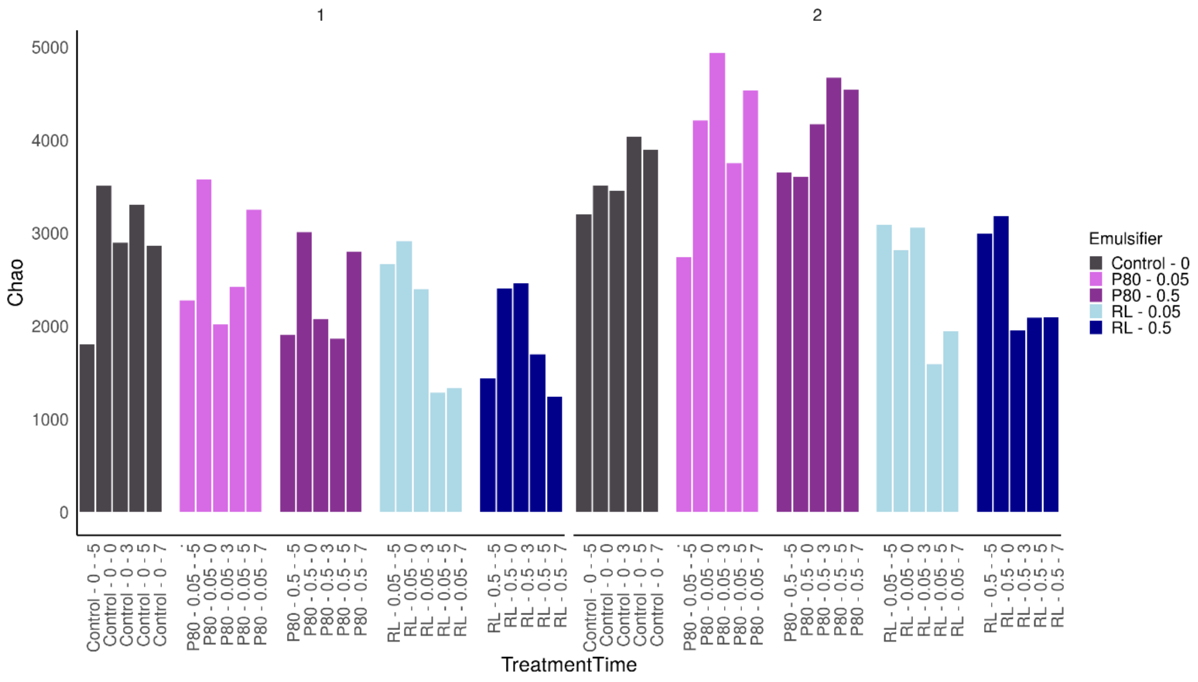


*Figure 5: Chao richness of the simulated gut microbiota in mucosal compartments of a 16 and 18 day SHIME experiment investigating the impact of TWEEN80 and rhamnolipids (0,05 m% and 0,5 m%) on the gut microbiota from two human faecal donors.*

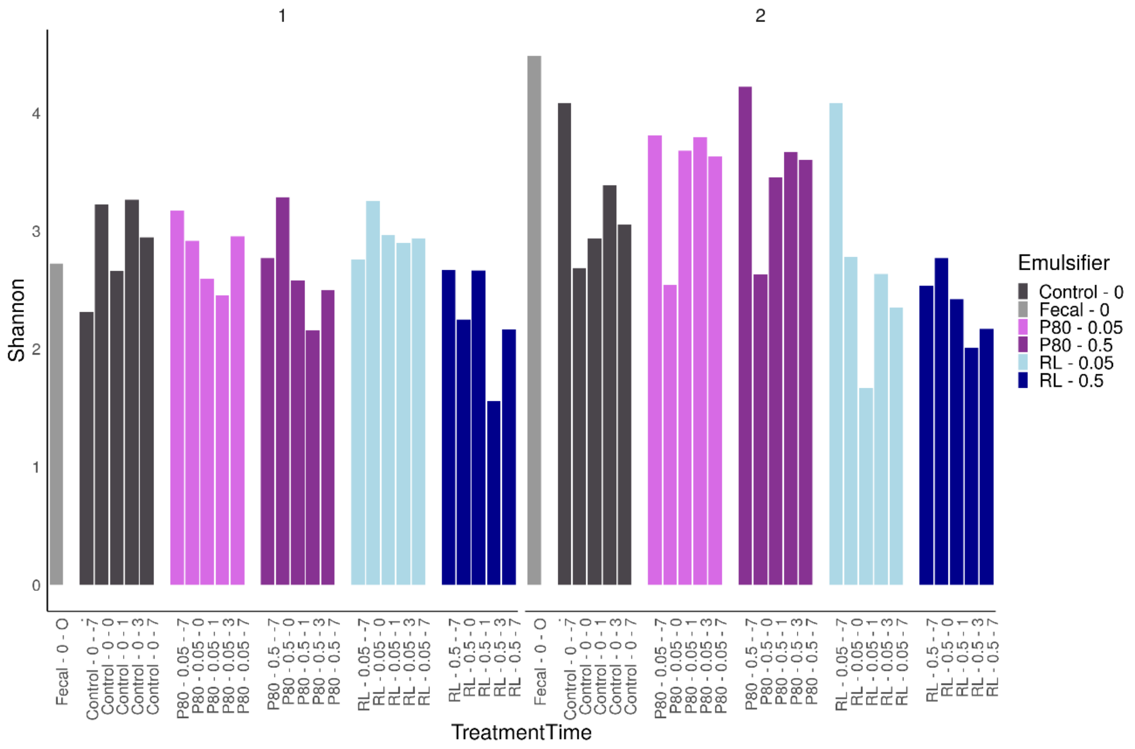


*Figure 6: Shannon index for the simulated gut microbiota in luminal suspensions from a 16 and 18 day SHIME experiment investigating the impact of TWEEN80 and rhamnolipids (0,05 m% and 0,5 m%) on the gut microbiota from two human faecal donors.*

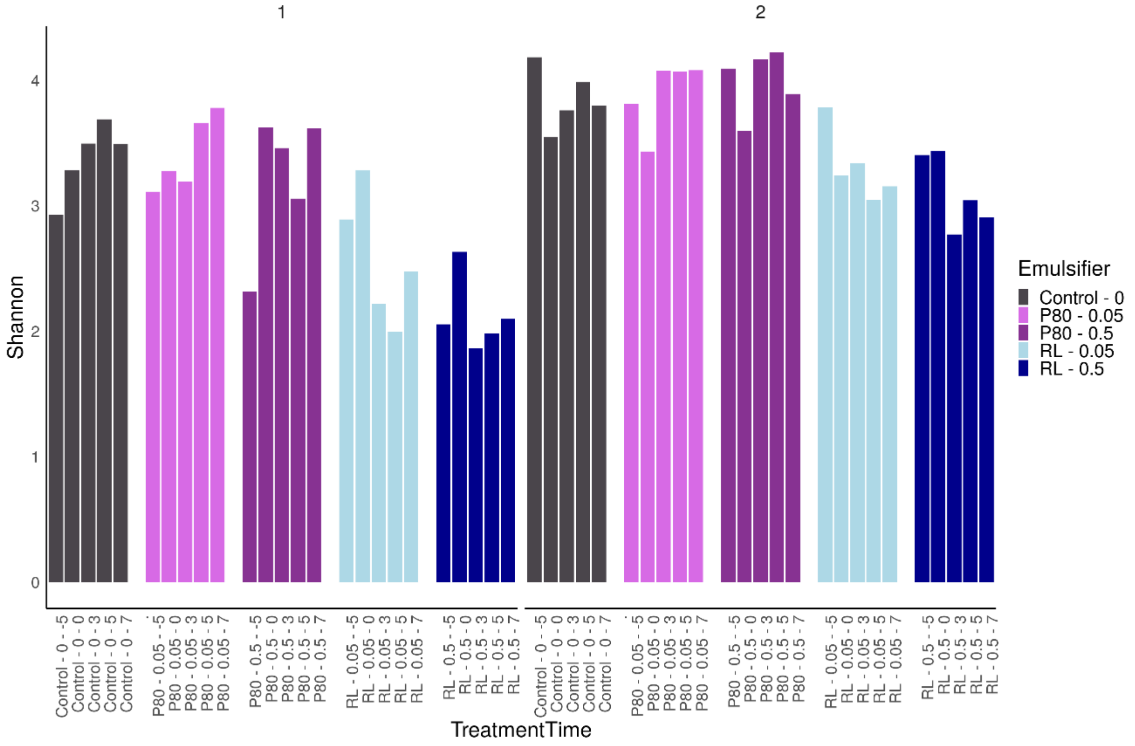


*Figure 7: Shannon index for the simulated gut microbiota in mucosal compartments of a 16 and 18 day SHIME experiment investigating the impact of TWEEN80 and rhamnolipids (0,05 m% and 0,5 m%) on the gut microbiota from two human faecal donors.*

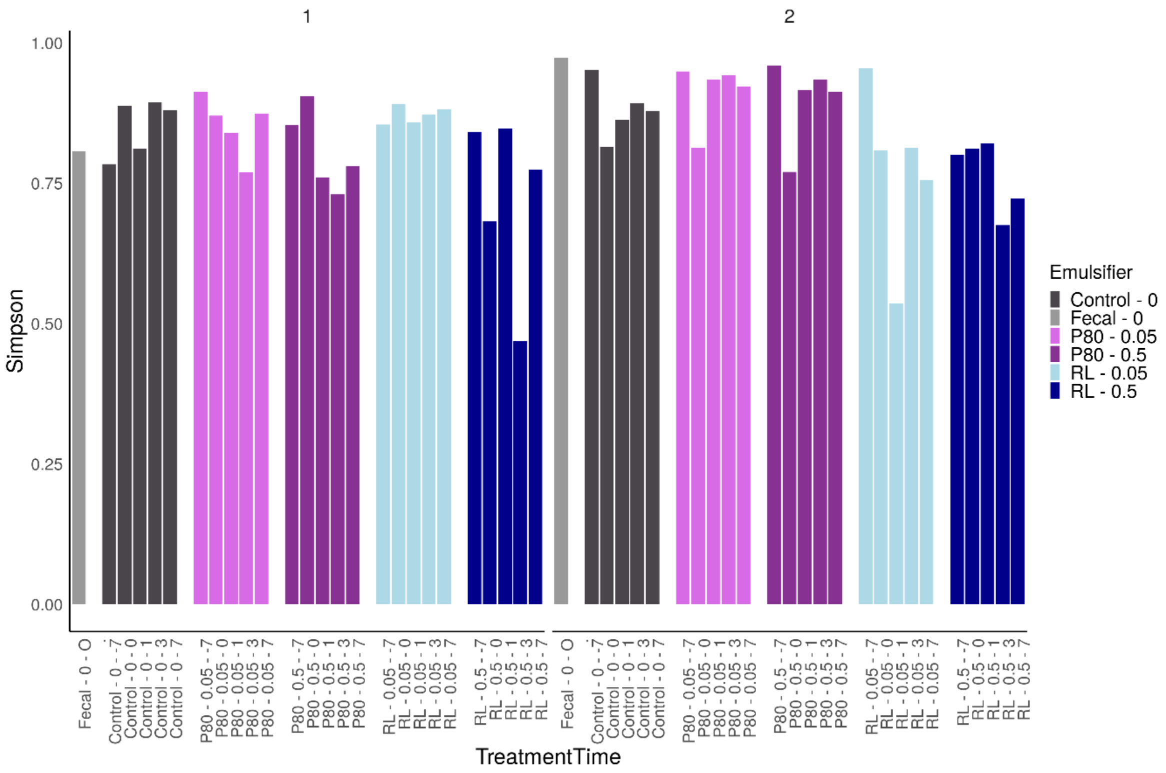


*Figure 8: Simpson index for the simulated gut microbiota in luminal suspensions from a 16 and 18 day SHIME experiment investigating the impact of TWEEN80 and rhamnolipids (0,05 m% and 0,5 m%) on the gut microbiota from two human faecal donors.*

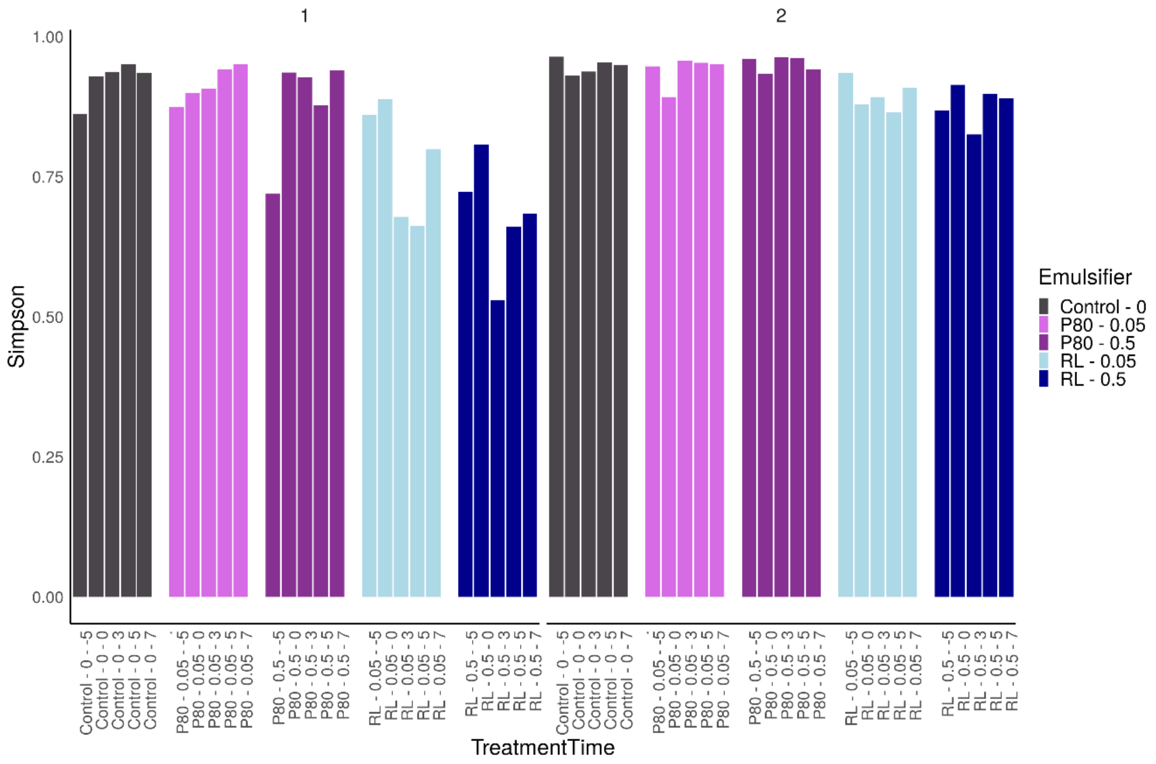


*Figure 9: Simpson index for the simulated gut microbiota in mucosal compartments of a 16 and 18 day SHIME experiment investigating the impact of TWEEN80 and rhamnolipids (0,05 m% and 0,5 m%) on the gut microbiota from two human faecal donors.*

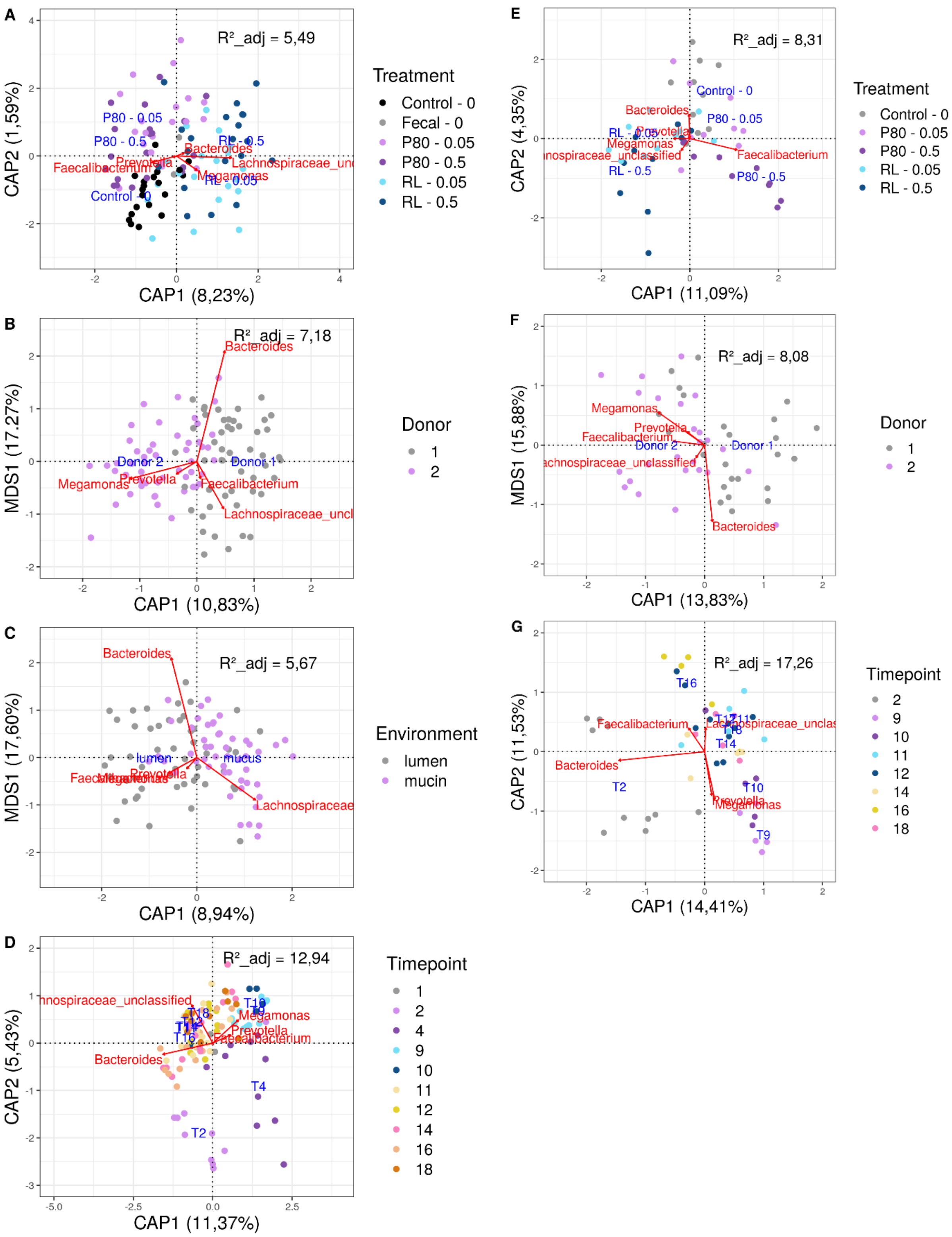


*Figure 10: Type II scaling triplot obtained from partial distance based redundancy analysis for the abundances of microbial genera in samples from a 16 and 18 day SHIME-esperiment investigating the impact of a 7 day treatment with TWEEN80 and rhamnolipids (0,05 m% and 0,5 m%) on the gut microbiota from two human faecal donors. Abundances were retrieved by use of 16S rRNA amplicon sequencing. Plots A, B, C and D visualise the variation comprised by the factors Treatment, Donor, Location and Timepoint in the relative abundance data. Plots E, F and G visualize the variation explained by factors Treatment, Donor and Timepoint in the absolute abundance data. In each plot the factor levels were set as explanatory variables (blue arrows) and the 5 most prevalent genera were set as response variables (red arrows).*

*Table 1: Log2FoldChange (L2FC) values and P-values of Wald test during DESeq-analysis of 16S rRNA Illumina amplicon sequencing data from luminal suspensions of a 16 and 18 day SHIME experiment investigating the impact of TWEEN80 and rhamnolipids (0,05 m% and 0,5 m%) on the gut microbiota from two human faecal donors.*

| LogFoldChange | | | | | P-values | | | |
|---|---|---|---|---|---|---|---|---|
| **P80_0.05** | **P80_0.5** | **RL_0.05** | **RL_0.5** | **Genus** | **P80_0.05** | **P80_0.5** | **RL_0.05** | **RL_0.5** |
| 0,02 | 0,14 | -4,82 | -4,86 | Agathobacter | 1,000 | 1,000 | **0,000** | **0,000** |
| 4,92 | 0,02 | 3,20 | 0,57 | Akkermansia | **0,000** | 1,000 | **0,010** | 0,425 |
| -1,52 | -3,12 | -2,53 | -5,46 | Alistipes | **0,040** | **0,000** | **0,002** | **0,000** |
| -0,26 | 0,48 | 1,43 | -3,31 | Anaeroglobus | 0,470 | 0,376 | 0,058 | **0,002** |
| 0,01 | 1,19 | -3,31 | -3,45 | Anaerostipes | 1,000 | 0,051 | **0,000** | **0,000** |
| -0,02 | -0,37 | -0,01 | -1,19 | Bacteroidales_unclassified | 1,000 | 0,368 | 1,000 | **0,029** |
| 0,00 | -1,51 | 0,00 | 0,07 | Bacteroides | 1,000 | **0,018** | 1,000 | 0,760 |
| 0,00 | -0,12 | -1,45 | -1,75 | Bifidobacteriaceae_unclassified | 1,000 | 0,988 | **0,048** | **0,029** |
| 0,04 | -1,12 | -5,71 | -4,52 | Bifidobacterium | 1,000 | 0,095 | **0,000** | **0,000** |
| 0,01 | -0,03 | -0,02 | 1,02 | Bilophila | 1,000 | 1,000 | 1,000 | **0,041** |
| 0,07 | 0,11 | -8,24 | -5,46 | Blautia | 1,000 | 1,000 | **0,000** | **0,000** |
| 0,03 | -0,02 | 0,19 | 2,32 | Butyricicoccaceae_unclassified | 1,000 | 1,000 | 0,635 | **0,011** |
| 0,04 | 0,29 | -0,66 | -1,27 | Butyricicoccus | 1,000 | 0,493 | 0,118 | **0,044** |
| 0,03 | 0,05 | -3,23 | -3,61 | Candidatus_Soleaferrea | 1,000 | 1,000 | **0,000** | **0,000** |
| -0,03 | -0,01 | -0,15 | -2,42 | Centipeda | 1,000 | 1,000 | 0,643 | **0,008** |
| 0,00 | 1,05 | -2,17 | -1,68 | Clostridia_unclassified | 1,000 | 0,051 | **0,007** | **0,016** |
| 0,00 | 0,11 | 0,00 | 4,63 | Clostridiaceae_unclassified | 1,000 | 1,000 | 1,000 | **0,000** |
| 0,00 | -1,22 | -3,28 | -3,70 | Colidextribacter | 1,000 | **0,045** | **0,000** | **0,000** |
| 1,59 | 3,05 | -3,53 | -3,27 | Collinsella | 0,051 | **0,001** | **0,002** | **0,002** |
| 0,92 | 0,21 | -2,79 | -2,81 | Coprococcus | 0,102 | 0,649 | **0,002** | **0,001** |
| 0,01 | 0,11 | -5,83 | -5,76 | Dorea | 1,000 | 1,000 | **0,000** | **0,000** |
| 0,70 | 2,34 | -1,32 | -0,68 | Eisenbergiella | 0,169 | **0,007** | 0,071 | 0,213 |
| 6,62 | 0,09 | 2,30 | 4,25 | Enterobacteriaceae_unclassified | **0,000** | 1,000 | **0,046** | **0,001** |
| 1,74 | 2,02 | -0,16 | -0,09 | Erysipelatoclostridium | **0,029** | **0,014** | 0,713 | 1,000 |
| -0,05 | 0,18 | -12,27 | -12,58 | Faecalibacterium | 1,000 | 0,649 | **0,000** | **0,000** |
| 0,03 | 0,05 | -3,72 | -3,60 | Fusicatenibacter | 1,000 | 1,000 | **0,000** | **0,000** |
| -0,67 | -2,11 | -0,13 | -4,22 | Gastranaerophilales_ge | 0,169 | **0,016** | 0,715 | **0,000** |
| 0,17 | -0,34 | -2,86 | -2,89 | Holdemanella | 0,554 | 0,493 | **0,003** | **0,002** |
| 0,06 | -0,03 | -1,40 | -1,60 | Holdemania | 1,000 | 1,000 | **0,046** | **0,030** |
| 2,33 | 1,72 | -0,78 | -0,23 | Hungatella | **0,003** | **0,019** | 0,109 | 0,460 |
| -0,30 | 0,08 | 0,03 | 3,26 | Lachnoclostridium | 0,468 | 1,000 | 1,000 | **0,003** |
| 0,14 | 1,40 | -2,94 | -2,98 | Lachnospira | 0,759 | 0,051 | **0,005** | **0,003** |
| -0,74 | -1,09 | -6,07 | -3,57 | Lachnospiraceae_ge | 0,169 | 0,102 | **0,000** | **0,000** |
| 0,02 | -0,01 | -4,24 | -4,22 | Lachnospiraceae_ND3007_group | 1,000 | 1,000 | **0,000** | **0,000** |
| 0,00 | 0,00 | 0,30 | 2,10 | Lachnospiraceae_unclassified | 1,000 | 1,000 | 0,124 | **0,000** |
| 0,13 | 0,02 | 0,02 | 2,04 | Lactobacillales_unclassified | 0,759 | 1,000 | 1,000 | **0,010** |
| -0,58 | -0,05 | -0,05 | 1,47 | Lactobacillus | 0,183 | 1,000 | 1,000 | **0,019** |
| 0,03 | -0,08 | -1,24 | -3,68 | Megasphaera | 1,000 | 1,000 | 0,067 | **0,000** |
| -0,02 | -0,04 | 0,91 | 2,79 | Micrococcaceae_unclassified | 1,000 | 1,000 | 0,098 | **0,001** |
| -0,02 | -1,67 | -3,53 | -3,75 | Negativibacillus | 1,000 | **0,019** | **0,000** | **0,000** |
| 0,42 | 2,54 | -0,12 | -0,12 | Oscillibacter | 0,231 | **0,000** | 0,772 | 0,879 |
| 1,06 | 1,83 | 0,00 | 0,00 | Oscillospira | 0,099 | **0,019** | 1,000 | 1,000 |
| 0,01 | -0,02 | -3,66 | -3,66 | Oscillospirales_unclassified | 1,000 | 1,000 | **0,000** | **0,000** |

| | | | | | | | | |
|---|---|---|---|---|---|---|---|---|
| -0,01 | -0,02 | 0,04 | -3,39 | Parabacteroides | 1,000 | 1,000 | 1,000 | **0,000** |
| -0,04 | -0,53 | -2,87 | -4,13 | Paraprevotella | 1,000 | 0,331 | **0,002** | **0,000** |
| -0,09 | -0,35 | -0,03 | -4,55 | Prevotella | 1,000 | 0,512 | 1,000 | **0,000** |
| -0,05 | -0,14 | -0,26 | -2,80 | Prevotellaceae_unclassified | 1,000 | 0,917 | 0,408 | **0,002** |
| 0,37 | 1,52 | -6,45 | -5,95 | Ruminococcaceae_unclassified | 0,298 | **0,044** | **0,000** | **0,000** |
| 3,15 | 0,14 | -3,60 | -3,38 | Subdoligranulum | **0,000** | 0,988 | **0,000** | **0,000** |
| -0,01 | -0,02 | -1,26 | -1,85 | Sutterella | 1,000 | 1,000 | **0,030** | **0,005** |
| -0,12 | 0,04 | -3,74 | -5,15 | Tyzzerella | 0,852 | 1,000 | **0,000** | **0,000** |
| 0,00 | -0,04 | -1,88 | -2,01 | UBA1819 | 1,000 | 1,000 | **0,018** | **0,012** |
| 2,18 | 0,17 | -0,02 | -0,03 | UCG-002 | **0,013** | 0,879 | 1,000 | 1,000 |
| -0,01 | 0,19 | -2,33 | -2,40 | UCG-003 | 1,000 | 0,649 | **0,007** | **0,005** |

*Table 2: Log2FoldChange (L2FC) values and P-values of Wald test during DESeq-analysis of 16S rRNA Illumina amplicon sequencing data from mucosal suspensions of a 16 and 18 day SHIME experiment investigating the impact of TWEEN80 and rhamnolipids (0,05 m% and 0,5 m%) on the gut microbiota from two human faecal donors.*

| LogFoldChange | | | | | P-values | | | |
|---|---|---|---|---|---|---|---|---|
| **P80_0.05** | **P80_0.5** | **RL_0.05** | **RL_0.5** | **Genus** | **P80_0.05** | **P80_0.5** | **RL_0.05** | **RL_0.5** |
| -0,01 | -0,12 | -2,40 | -2,82 | Acetanaerobacterium | 1,000 | 0,896 | 0,001 | 0,000 |
| 0,01 | 0,09 | -3,23 | -5,16 | Agathobacter | 1,000 | 1,000 | 0,000 | 0,000 |
| 4,69 | 0,03 | 3,87 | 0,22 | Akkermansia | 0,000 | 1,000 | 0,001 | 0,970 |
| -0,02 | -0,14 | -0,94 | -5,12 | Alistipes | 1,000 | 0,816 | 0,054 | 0,000 |
| -3,01 | 0,09 | 0,45 | -5,43 | Anaeroglobus | 0,009 | 1,000 | 0,476 | 0,000 |
| -0,02 | 0,06 | -3,36 | -3,81 | Anaerostipes | 1,000 | 1,000 | 0,000 | 0,000 |
| 2,54 | 0,22 | -0,50 | -0,98 | Barnesiella | 0,002 | 0,596 | 0,258 | 0,104 |
| 0,00 | -0,36 | -3,17 | -4,17 | Bifidobacterium | 1,000 | 0,460 | 0,001 | 0,000 |
| -0,03 | -0,65 | -2,14 | -0,99 | Bilophila | 1,000 | 0,142 | 0,000 | 0,027 |
| -0,03 | -0,09 | -5,64 | -6,48 | Blautia | 1,000 | 1,000 | 0,000 | 0,000 |
| 0,03 | 0,06 | -1,18 | -2,41 | CAG-56 | 1,000 | 1,000 | 0,068 | 0,004 |
| -0,01 | 0,10 | -3,02 | -3,60 | Candidatus_Soleaferrea | 1,000 | 1,000 | 0,000 | 0,000 |
| 0,00 | 0,50 | -0,44 | -4,31 | Centipeda | 1,000 | 0,299 | 0,295 | 0,000 |
| -1,09 | -1,62 | -0,17 | -0,18 | Cloacibacillus | 0,066 | 0,034 | 0,563 | 0,530 |
| 0,00 | -0,03 | -0,34 | -1,71 | Clostridia_unclassified | 1,000 | 1,000 | 0,258 | 0,014 |
| -0,01 | -2,51 | -4,43 | -4,91 | Colidextribacter | 1,000 | 0,000 | 0,000 | 0,000 |
| 0,01 | 0,02 | -3,13 | -3,48 | Collinsella | 1,000 | 1,000 | 0,000 | 0,000 |
| 0,10 | 0,17 | -2,57 | -3,15 | Coprococcus | 0,875 | 0,715 | 0,002 | 0,000 |
| 0,00 | -0,16 | -0,76 | -3,50 | Desulfovibrio | 1,000 | 0,729 | 0,124 | 0,000 |
| 0,03 | -0,23 | -0,94 | -3,37 | Desulfovibrionaceae_unclassified | 1,000 | 0,596 | 0,113 | 0,000 |
| -0,01 | 0,07 | -5,40 | -6,43 | Dorea | 1,000 | 1,000 | 0,000 | 0,000 |
| 0,11 | 0,30 | -2,87 | -3,55 | Eisenbergiella | 0,841 | 0,516 | 0,001 | 0,000 |
| 0,06 | 0,05 | 0,98 | 3,55 | Enterobacterales_unclassified | 1,000 | 1,000 | 0,115 | 0,000 |
| 4,24 | 0,13 | 6,07 | 6,58 | Enterobacteriaceae_unclassified | 0,001 | 1,000 | 0,000 | 0,000 |
| 0,28 | -0,17 | -1,89 | -3,16 | Erysipelotrichaceae_ge | 0,337 | 0,710 | 0,011 | 0,000 |
| -0,20 | 0,06 | -6,89 | -9,24 | Faecalibacterium | 0,518 | 1,000 | 0,000 | 0,000 |
| 0,04 | 0,55 | -0,32 | -3,32 | Flavonifractor | 1,000 | 0,255 | 0,376 | 0,000 |
| 0,03 | 0,24 | -2,56 | -3,64 | Fusicatenibacter | 1,000 | 0,595 | 0,002 | 0,000 |
| -0,03 | 0,06 | 0,05 | -3,44 | Gastranaerophilales_ge | 1,000 | 1,000 | 1,000 | 0,000 |
| -0,06 | -0,88 | -0,59 | -2,51 | Holdemanella | 1,000 | 0,142 | 0,227 | 0,004 |
| -0,06 | -1,30 | -2,40 | -2,97 | Holdemania | 1,000 | 0,072 | 0,002 | 0,000 |
| 1,34 | 0,43 | -1,01 | -3,65 | Hungatella | 0,040 | 0,349 | 0,068 | 0,000 |
| 0,91 | 0,57 | -0,69 | -1,75 | Intestinimonas | 0,103 | 0,255 | 0,204 | 0,027 |
| -0,02 | 0,06 | -3,60 | -4,49 | Lachnospira | 1,000 | 1,000 | 0,000 | 0,000 |
| -2,91 | -2,66 | -5,44 | -4,14 | Lachnospiraceae_ge | 0,002 | 0,000 | 0,000 | 0,000 |
| 0,01 | 0,10 | -1,49 | -3,06 | Lachnospiraceae_ND3007_group | 1,000 | 1,000 | 0,047 | 0,000 |
| 0,00 | -0,19 | 1,21 | 1,37 | Lachnospiraceae_unclassified | 1,000 | 0,408 | 0,000 | 0,000 |
| 0,21 | 0,00 | -0,01 | 1,92 | Lactobacillales_unclassified | 0,470 | 1,000 | 1,000 | 0,009 |
| -1,68 | -0,09 | -0,56 | -0,16 | Lactobacillus | 0,036 | 1,000 | 0,204 | 0,676 |
| 0,00 | -0,02 | -2,68 | -2,71 | Marvinbryantia | 1,000 | 1,000 | 0,000 | 0,000 |
| -0,25 | -0,14 | -0,04 | -1,18 | Megamonas | 0,337 | 0,820 | 1,000 | 0,038 |
| 0,03 | -0,09 | -2,38 | -4,70 | Megasphaera | 1,000 | 1,000 | 0,006 | 0,000 |
| -0,04 | -1,23 | -3,44 | -4,32 | Negativibacillus | 1,000 | 0,072 | 0,000 | 0,000 |

| | | | | | | | | |
|---|---|---|---|---|---|---|---|---|
| 0,61 | 0,05 | -0,16 | -2,51 | Odoribacter | 0,147 | 1,000 | 0,641 | 0,001 |
| 0,01 | 0,11 | -3,00 | -3,59 | Oscillibacter | 1,000 | 0,933 | 0,000 | 0,000 |
| 1,30 | 0,92 | -1,50 | -2,47 | Oscillospira | 0,039 | 0,099 | 0,035 | 0,004 |
| 0,04 | -0,09 | -3,65 | -5,06 | Oscillospiraceae_unclassified | 1,000 | 1,000 | 0,000 | 0,000 |
| 0,02 | -0,22 | -1,55 | -2,21 | Oscillospirales_unclassified | 1,000 | 0,595 | 0,020 | 0,003 |
| 0,05 | 0,09 | -0,02 | -3,90 | Parabacteroides | 1,000 | 1,000 | 1,000 | 0,000 |
| -0,21 | -0,37 | -2,72 | -5,43 | Paraprevotella | 0,470 | 0,408 | 0,001 | 0,000 |
| 0,01 | 0,00 | -2,00 | -2,44 | Phocea | 1,000 | 1,000 | 0,007 | 0,001 |
| -0,31 | 0,03 | 0,01 | -5,42 | Prevotella | 0,337 | 1,000 | 1,000 | 0,000 |
| -1,51 | -0,18 | -0,30 | -2,93 | Prevotellaceae_unclassified | 0,036 | 0,657 | 0,282 | 0,000 |
| -0,11 | -0,22 | 1,78 | 2,45 | Pseudomonas | 0,530 | 0,595 | 0,001 | 0,000 |
| 0,03 | 0,23 | -4,30 | -4,88 | Ruminococcaceae_unclassified | 1,000 | 0,602 | 0,000 | 0,000 |
| -0,04 | -0,05 | -0,06 | -1,60 | Selenomonadaceae_unclassified | 1,000 | 1,000 | 1,000 | 0,021 |
| 0,02 | -0,62 | -3,85 | -4,66 | Subdoligranulum | 1,000 | 0,253 | 0,000 | 0,000 |
| -0,02 | 0,08 | -3,15 | -3,57 | Tyzzerella | 1,000 | 1,000 | 0,000 | 0,000 |
| -0,02 | -0,07 | -1,45 | -1,98 | UBA1819 | 1,000 | 1,000 | 0,033 | 0,009 |
| 0,23 | -0,09 | -1,75 | -2,25 | UCG-002 | 0,337 | 1,000 | 0,018 | 0,004 |
| -0,03 | -0,05 | -3,50 | -4,43 | UCG-003 | 1,000 | 1,000 | 0,000 | 0,000 |
| -0,01 | 0,22 | -0,07 | 4,25 | uncultured_ge | 1,000 | 0,646 | 1,000 | 0,000 |

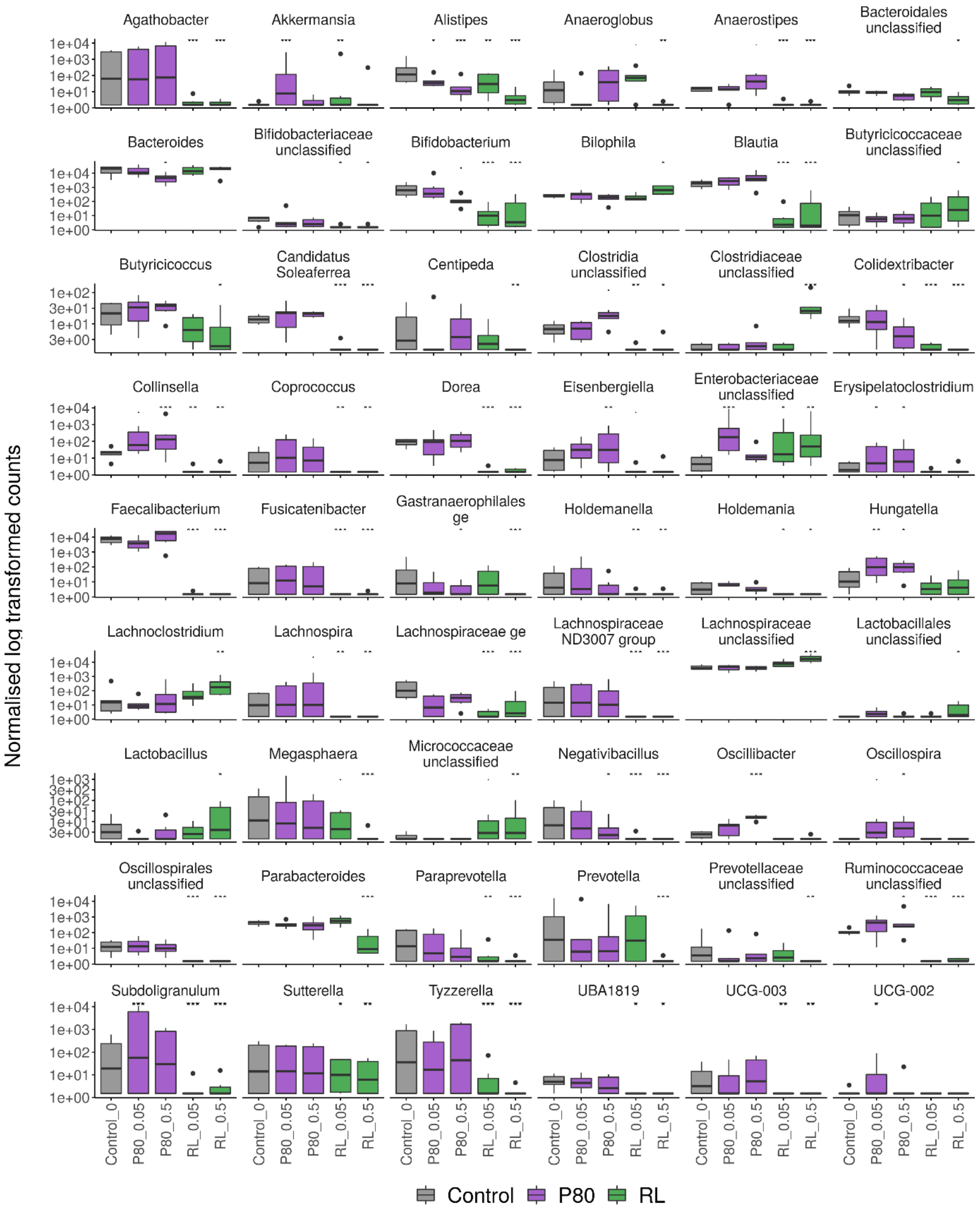


*Figure 11: Normalized counts of significantly increased or decreased genera, obtained through DESeq analysis of 16S rRNA Illumina amplicon sequencing data from luminal compartments in a 16 and 18 day SHIME experiment investigating the impact of TWEEN80 and rhamnolipids (0,05 m% and 0,5 m%) on the gut microbiota from two human faecal donors. Asterisks represent significant differences with the control based on a Wald test (α = 0,05).*

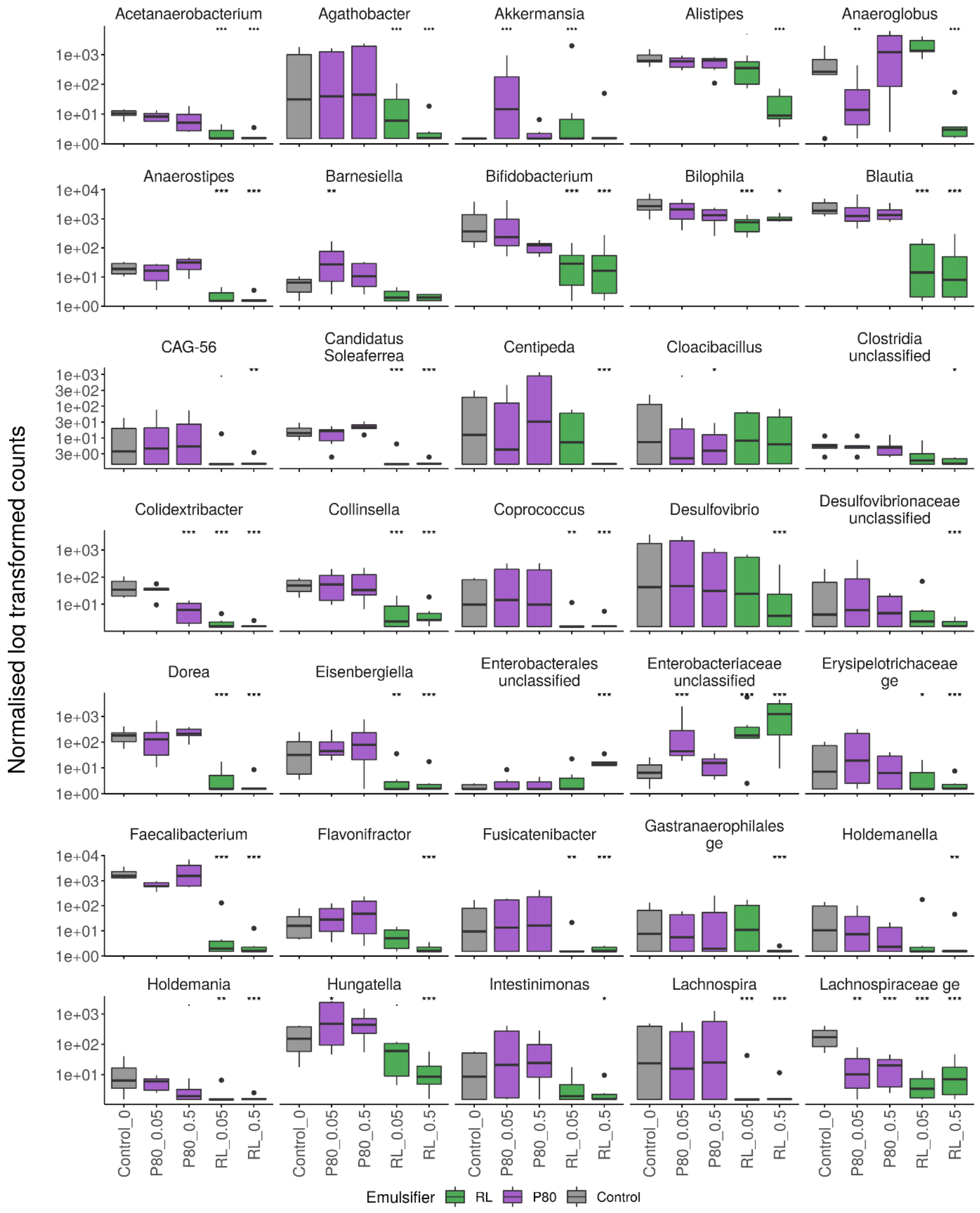
Normalised log transformed counts
Acetanaerobacterium
Agathobacter
Akkermansia
Alistipes
Anaeroglobus
Anaerostipes
Barnesiella
Bifidobacterium
Bilophila
Blautia
CAG-56
Candidatus Soleaferrea
Centipeda
Cloacibacillus
Clostridia unclassified
Colidextribacter
Collinsella
Coprococcus
Desulfovibrio
Desulfovibrionaceae unclassified
Dorea
Eisenbergiella
Enterobacterales unclassified
Enterobacteriaceae unclassified
Erysipelotrichaceae ge
Faecalibacterium
Flavonifractor
Fusicatenibacter
Gastranaerophilales ge
Holdemanella
Holdemania
Hungatella
Intestinimonas
Lachnospira
Lachnospiraceae ge
Control_0
P80_0.05
P80_0.5
RL_0.05
RL_0.5
Emulsifier
RL
P80
Control

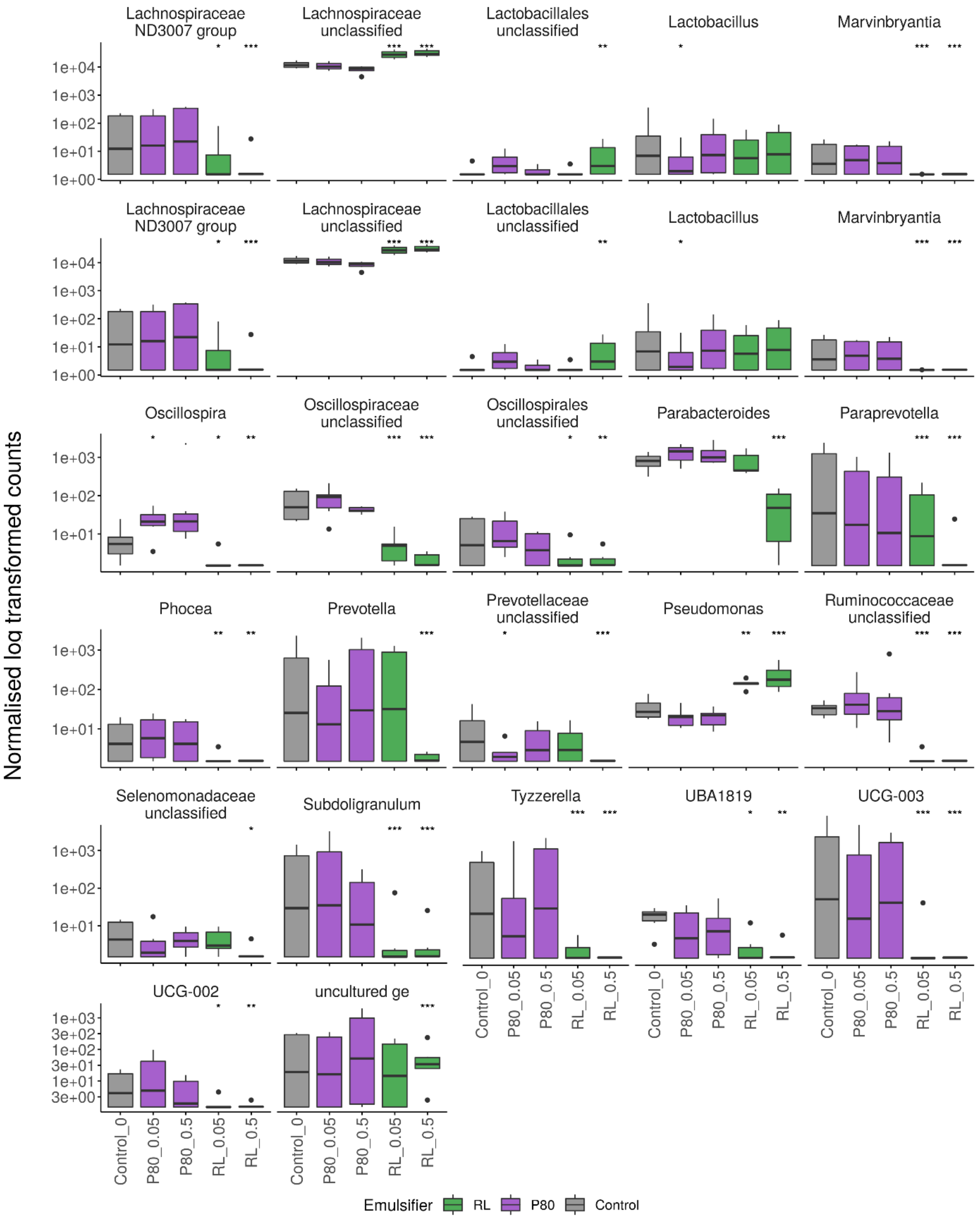


*Figure 12: Normalized counts of significantly increased or decreased genera, obtained through DESeq analysis of 16S rRNA Illumina amplicon sequencing data from mucosal compartments in a 16 and 18 day SHIME experiment investigating the impact of TWEEN80 and rhamnolipids (0,05 m% and 0,5 m%) on the gut microbiota from two human faecal donors. Asterisks represent significant differences with the control based on a Wald test (α = 0,05).*

#### 3. SCFA

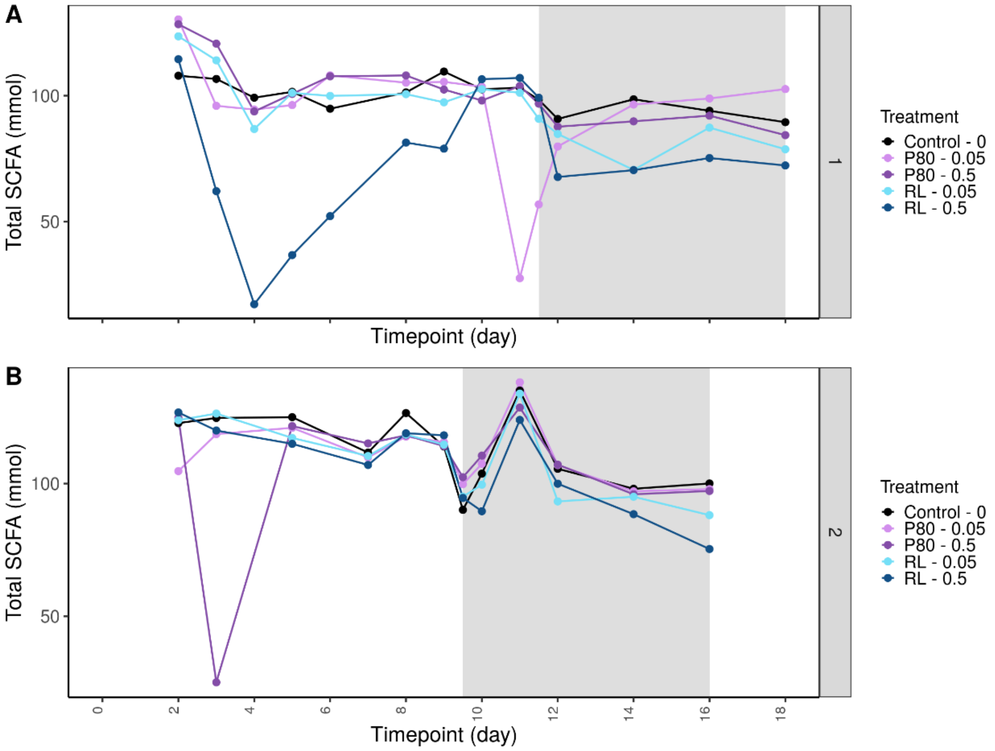


*Figure 13: Total SCFA levels (mM) in luminal suspension during a 16 and 18 day SHIME experiment investigating the impact of TWEEN80 and rhamnolipids (0,05 m% and 0,5 m%) on the gut microbiota of two human faecal donors. The 7-day treatment period is indicated by the grey background.*

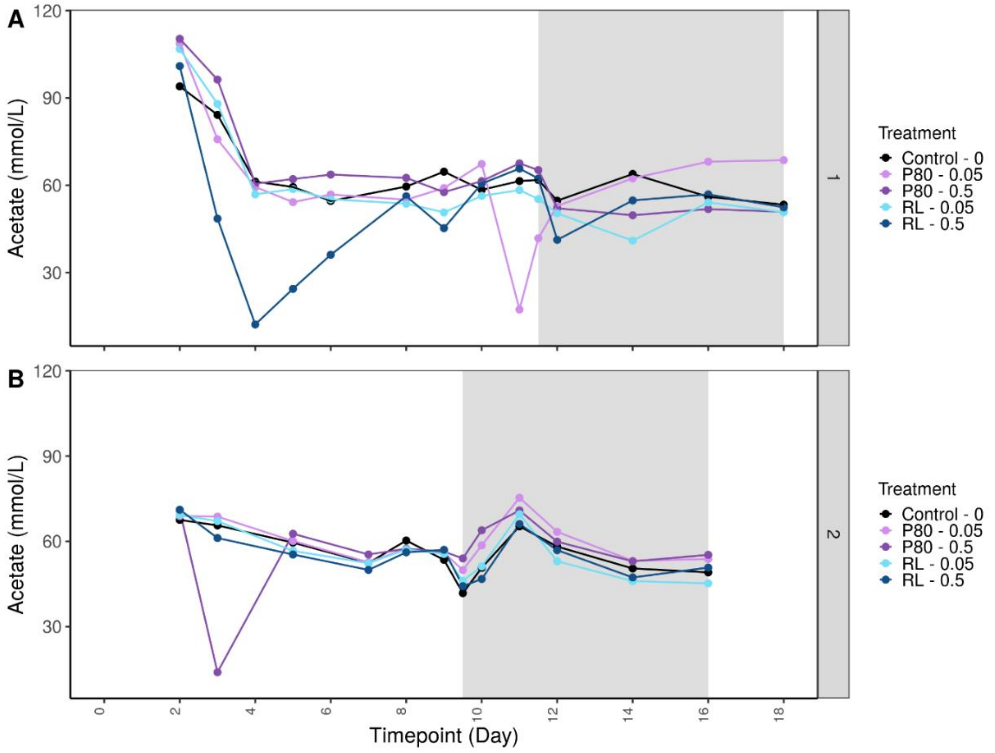


*Figure 14: Acetate levels (mM) in luminal suspension during a 16 and 18 day SHIME experiment investigating the impact of TWEEN80 and rhamnolipids (0,05 m% and 0,5 m%) on the gut microbiota of two human faecal donors. The 7-day treatment period is indicated by the grey background.*

### 4. Metabolomics

#### 4.1 Targeted

### 4.1.1 P80 – 0,05 m%

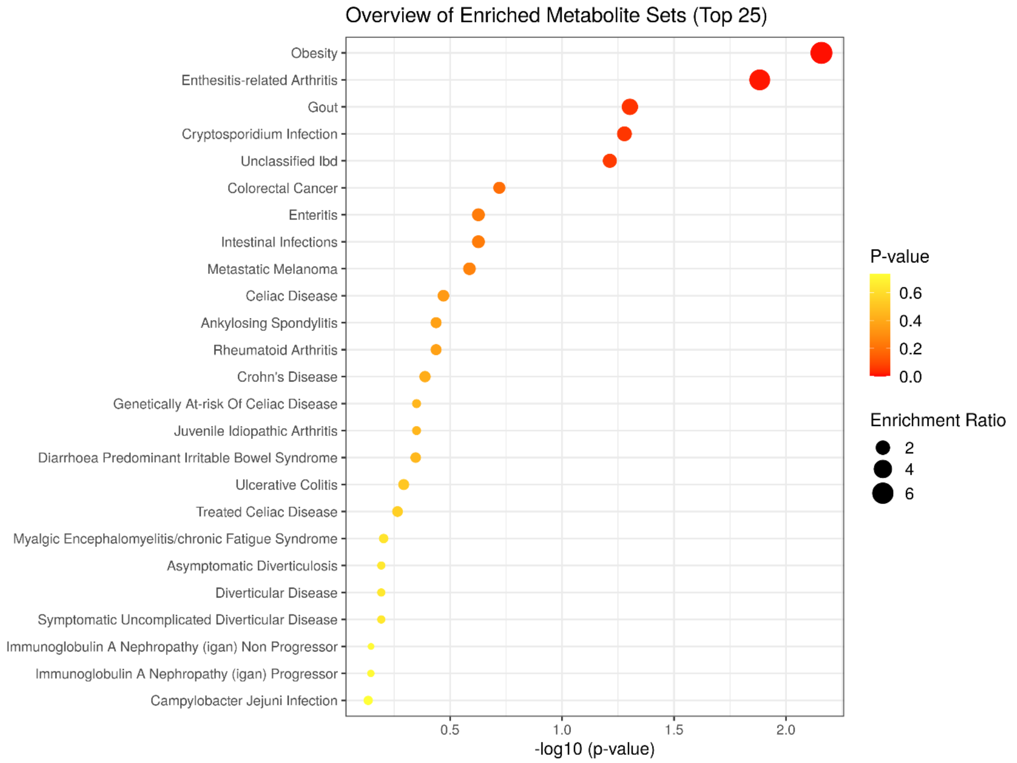


*Figure 15: Dot plot ranking disease features enriched in the human gut microbiota from two human faecal donors after exposure to 0,05% TWEEN80 for 7 days in the Simulator of the Human Intestinal Microbial Ecosystem (SHIME). Graph was obtained using the enrichment analysis tool on the MetaboAnalyst website.*

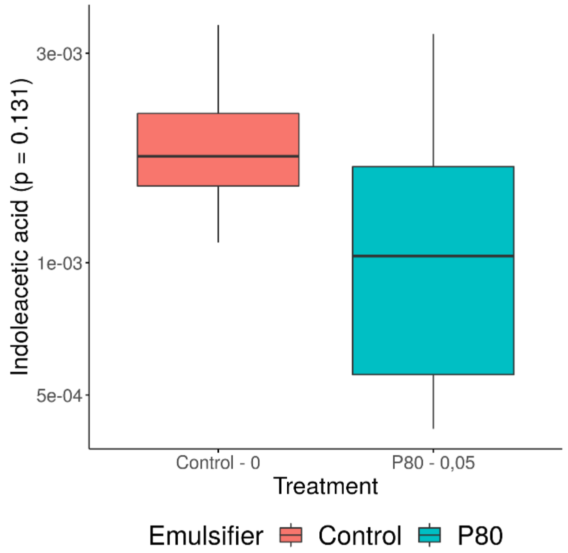


*Figure 16: Metabolites from the targeted metabolomics database annotated by the Enrichment Analysis tool on MetaboAnalyst as pointing towards Enthesitis-related Arthritis for the treatment with 0,05% of TWEEN80. Metabolites were detected in luminal suspensions from a 16 and 18 day SHIME experiment investigating the impact of TWEEN80 and rhamnolipids (0,05 m% and 0,5 m%) on the gut microbiota of two human faecal donors during a 7 day treatment period. Samples were from timepoint 2, 11, 12, 14 and 18 from SHIME 1 and timepoint 2, 9, 10, 12 and 16 from SHIME 2.*

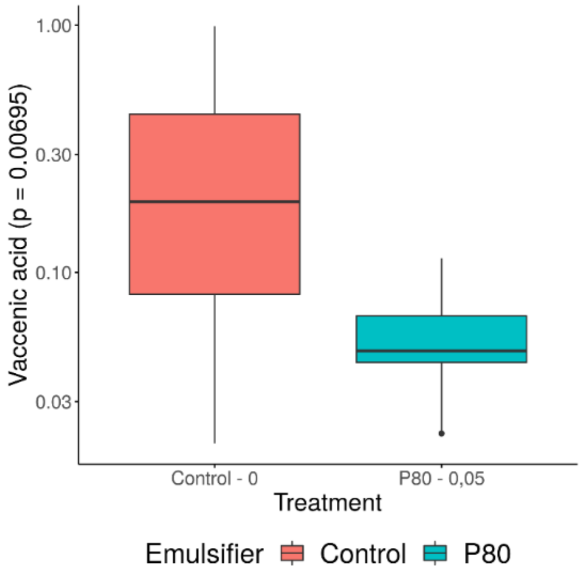


*Figure 17: Metabolites from the targeted metabolomics database annotated by the Enrichment Analysis tool on MetaboAnalyst as pointing towards Obesity for the treatment with 0,05% of TWEEN80. Metabolites were detected in luminal suspensions from a 16 and 18 day SHIME experiment investigating the impact of TWEEN80 and rhamnolipids (0,05 m% and 0,5 m%) on the gut microbiota of two human faecal donors during a 7 day treatment period. Samples were from timepoint 2, 11, 12, 14 and 18 from SHIME 1 and timepoint 2, 9, 10, 12 and 16 from SHIME 2.*

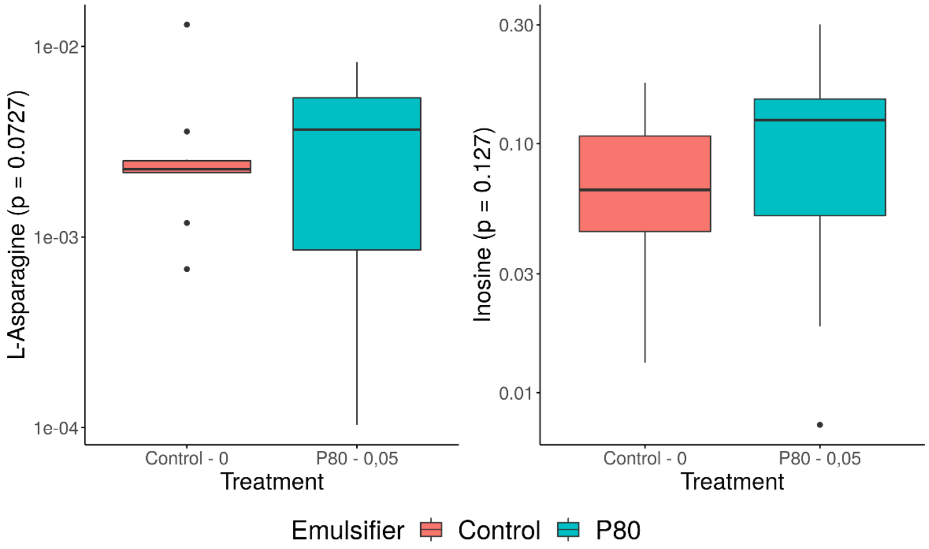


*Figure 18: Metabolites from the targeted metabolomics database annotated by the Enrichment Analysis tool on MetaboAnalyst as pointing towards Gout for the treatment with 0,05% of TWEEN80. Metabolites were detected in luminal suspensions from a 16 and 18 day SHIME experiment investigating the impact of TWEEN80 and rhamnolipids (0,05 m% and 0,5 m%) on the gut microbiota of two human faecal donors during a 7 day treatment period. Samples were from timepoint 2, 11, 12, 14 and 18 from SHIME 1 and timepoint 2, 9, 10, 12 and 16 from SHIME 2.*

### 4.1.2 P80 – 0,5 m%

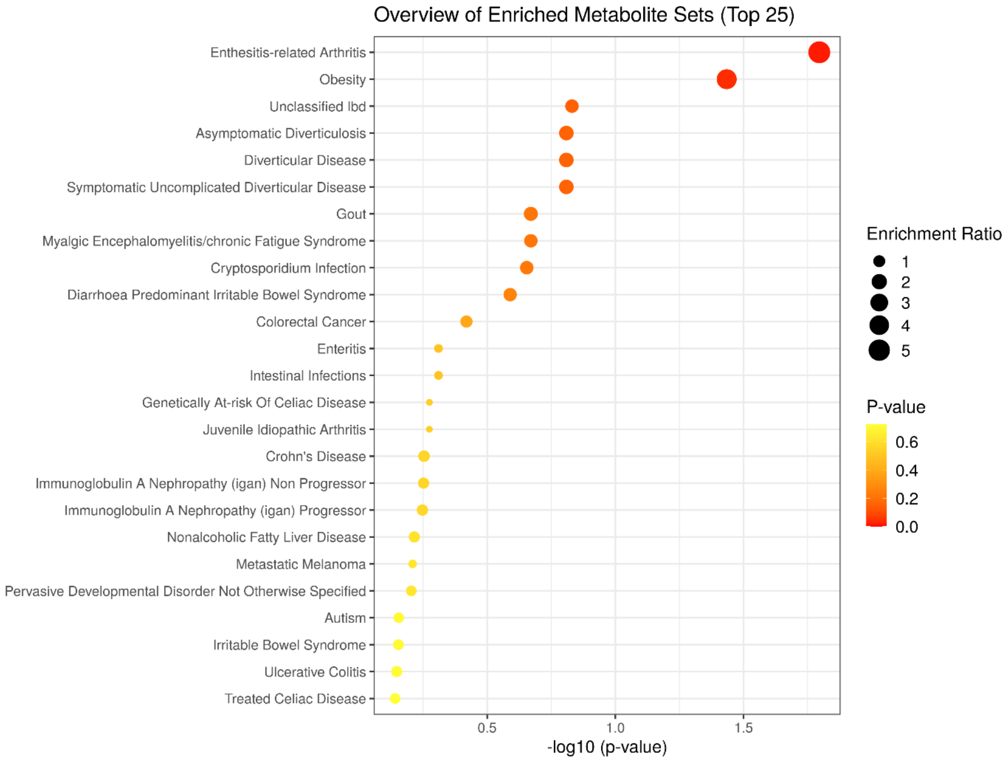


*Figure 19: Dot plot ranking disease features enriched in the human gut microbiota from two human faecal donors after exposure to 0,5% TWEEN80 for 7 days in the Simulator of the Human Intestinal Microbial Ecosystem (SHIME). Graph was obtained using the enrichment analysis tool on the MetaboAnalyst website.*

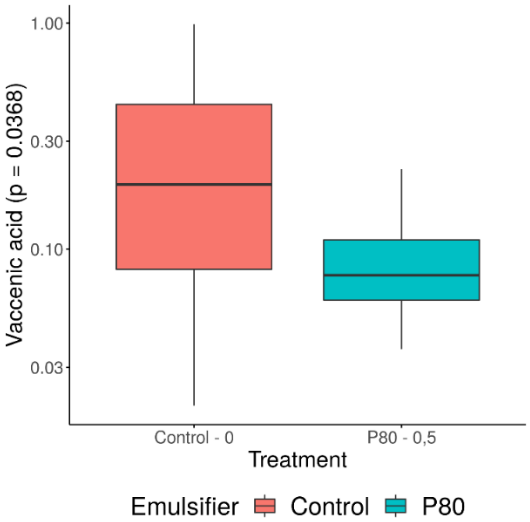


*Figure 20: Metabolites from the targeted metabolomics database annotated by the Enrichment Analysis tool on MetaboAnalyst as pointing towards Obesity for the treatment with 0,5% of TWEEN80. Metabolites were detected in luminal suspensions from a 16 and 18 day SHIME experiment investigating the impact of TWEEN80 and rhamnolipids (0,05 m% and 0,5 m%) on the gut microbiota of two human faecal donors during a 7 day treatment period. Samples were from timepoint 2, 11, 12, 14 and 18 from SHIME 1 and timepoint 2, 9, 10, 12 and 16 from SHIME 2.*

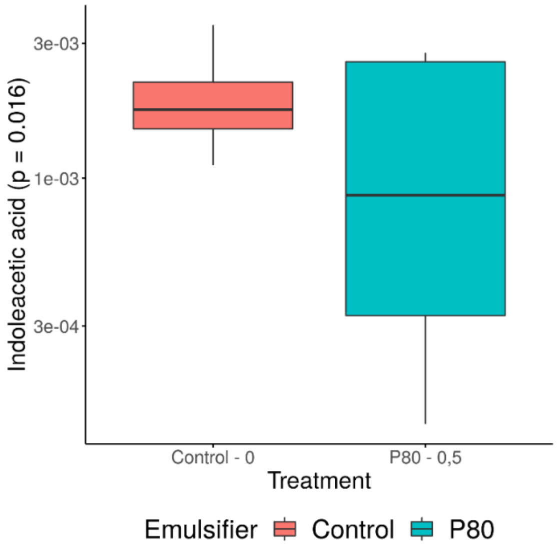


*Figure 21: Metabolites from the targeted metabolomics database annotated by the Enrichment Analysis tool on MetaboAnalyst as pointing towards Enthesitis-related Arthritis for the treatment with 0,5% of TWEEN80. Metabolites were detected in luminal suspensions from a 16 and 18 day SHIME experiment investigating the impact of TWEEN80 and rhamnolipids (0,05 m% and 0,5 m%) on the gut microbiota of two human faecal donors during a 7 day treatment period. Samples were from timepoint 2, 11, 12, 14 and 18 from SHIME 1 and timepoint 2, 9, 10, 12 and 16 from SHIME 2.*

### 4.1.3 RL – 0,05 m%

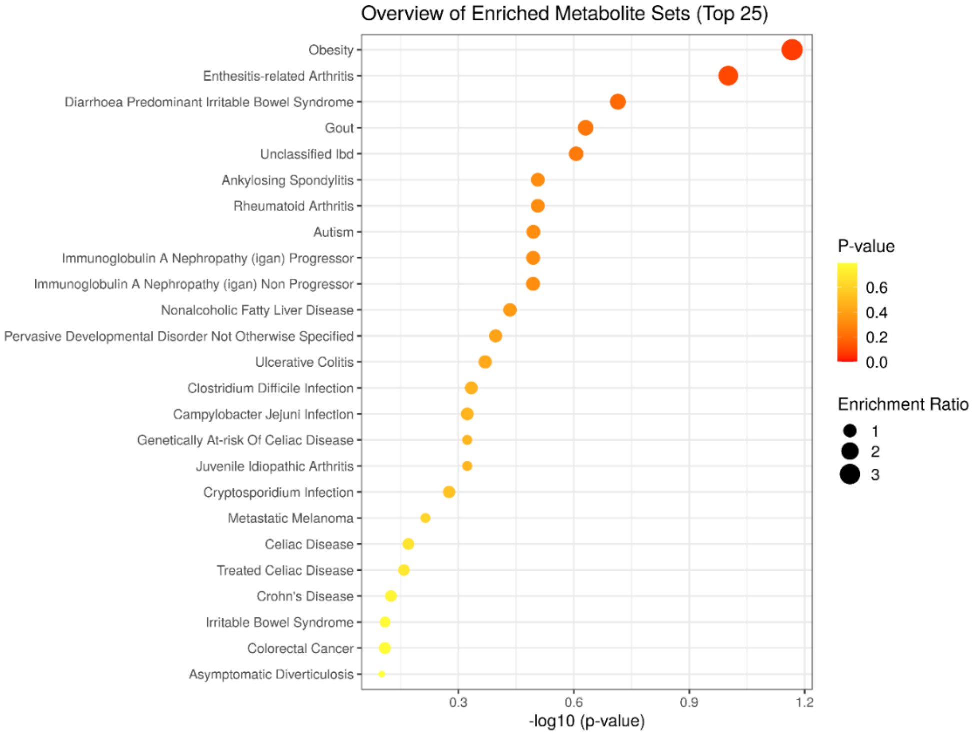


*Figure 22: Dot plot ranking disease features enriched in the human gut microbiota from two human faecal donors after exposure to 0,05% rhamnolipids for 7 days in the Simulator of the Human Intestinal Microbial Ecosystem (SHIME). Graph was obtained using the enrichment analysis tool on the MetaboAnalyst website.*

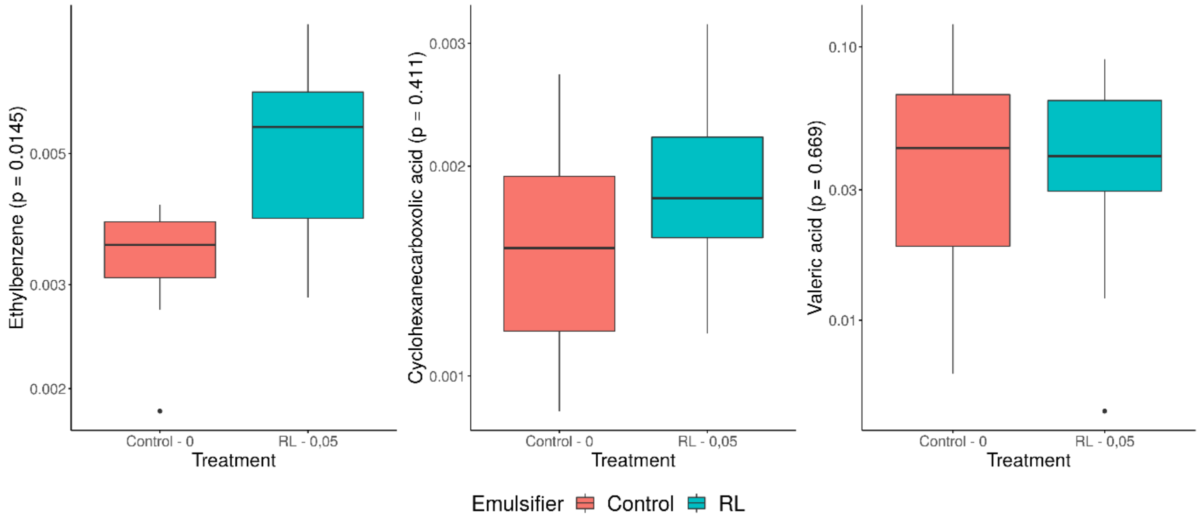


*Figure 23: Metabolites from the targeted metabolomics database annotated by the Enrichment Analysis tool on MetaboAnalyst as pointing towards Diarhea predominant IBS for the treatment with 0,05% of rhamnolipids. Metabolites were detected in luminal suspensions from a 16 and 18 day SHIME experiment investigating the impact of TWEEN80 and rhamnolipids (0,05 m% and 0,5 m%) on the gut microbiota of two human faecal donors during a 7 day treatment period. Samples were from timepoint 2, 11, 12, 14 and 18 from SHIME 1 and timepoint 2, 9, 10, 12 and 16 from SHIME 2.*

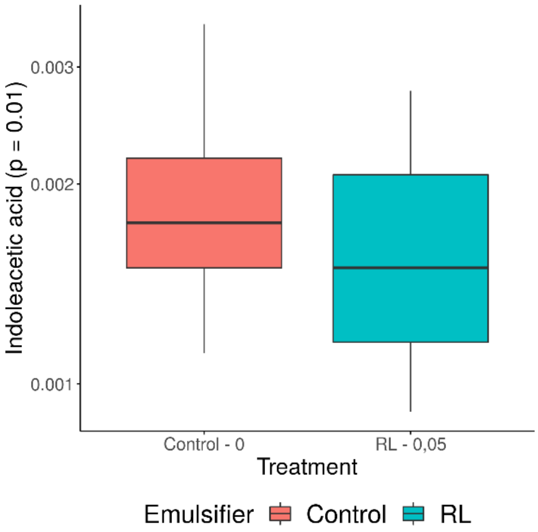


*Figure 24: Metabolites from the targeted metabolomics database annotated by the Enrichment Analysis tool on MetaboAnalyst as pointing towards Enthesitis related Arthritis for the treatment with 0,05% of rhamnolipids. Metabolites were detected in luminal suspensions from a 16 and 18 day SHIME experiment investigating the impact of TWEEN80 and rhamnolipids (0,05 m% and 0,5 m%) on the gut microbiota of two human faecal donors during a 7 day treatment period. Samples were from timepoint 2, 11, 12, 14 and 18 from SHIME 1 and timepoint 2, 9, 10, 12 and 16 from SHIME 2.*

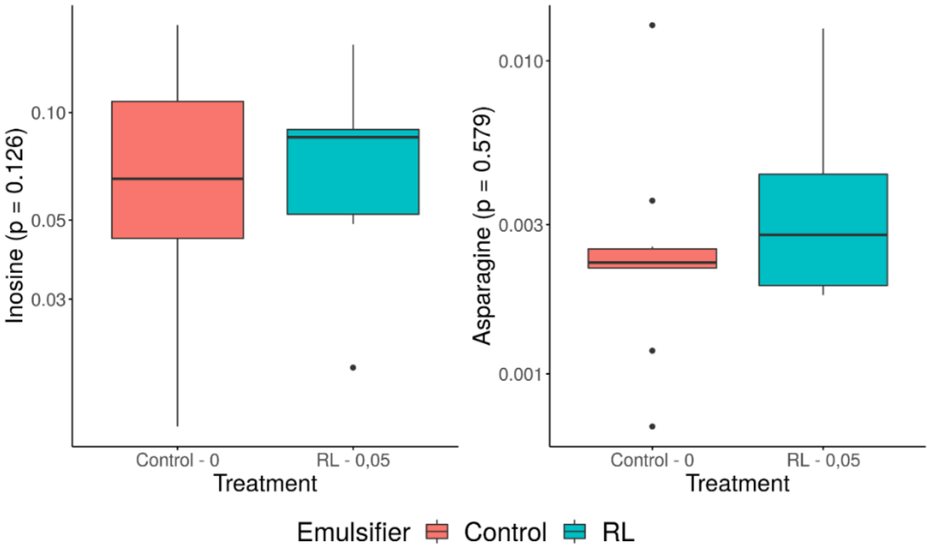


*Figure 25: Metabolites from the targeted metabolomics database annotated by the Enrichment Analysis tool on MetaboAnalyst as pointing towards Gout for the treatment with 0,05% of rhamnolipids. Metabolites were detected in luminal suspensions from a 16 and 18 day SHIME experiment investigating the impact of TWEEN80 and rhamnolipids (0,05 m% and 0,5 m%) on the gut microbiota of two human faecal donors during a 7 day treatment period. Samples were from timepoint 2, 11, 12, 14 and 18 from SHIME 1 and timepoint 2, 9, 10, 12 and 16 from SHIME 2.*

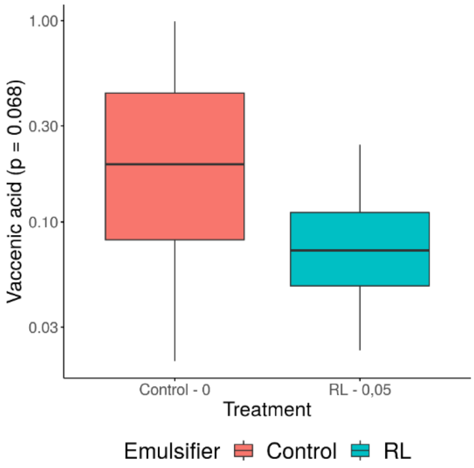


*Figure 26: Metabolites from the targeted metabolomics database annotated by the Enrichment Analysis tool on MetaboAnalyst as pointing towards Obesity for the treatment with 0,05% of rhamnolipids. Metabolites were detected in luminal suspensions from a 16 and 18 day SHIME experiment investigating the impact of TWEEN80 and rhamnolipids (0,05 m% and 0,5 m%) on the gut microbiota of two human faecal donors during a 7 day treatment period. Samples were from timepoint 2, 11, 12, 14 and 18 from SHIME 1 and timepoint 2, 9, 10, 12 and 16 from SHIME 2.*

### 4.1.4 RL – 0,5 m%

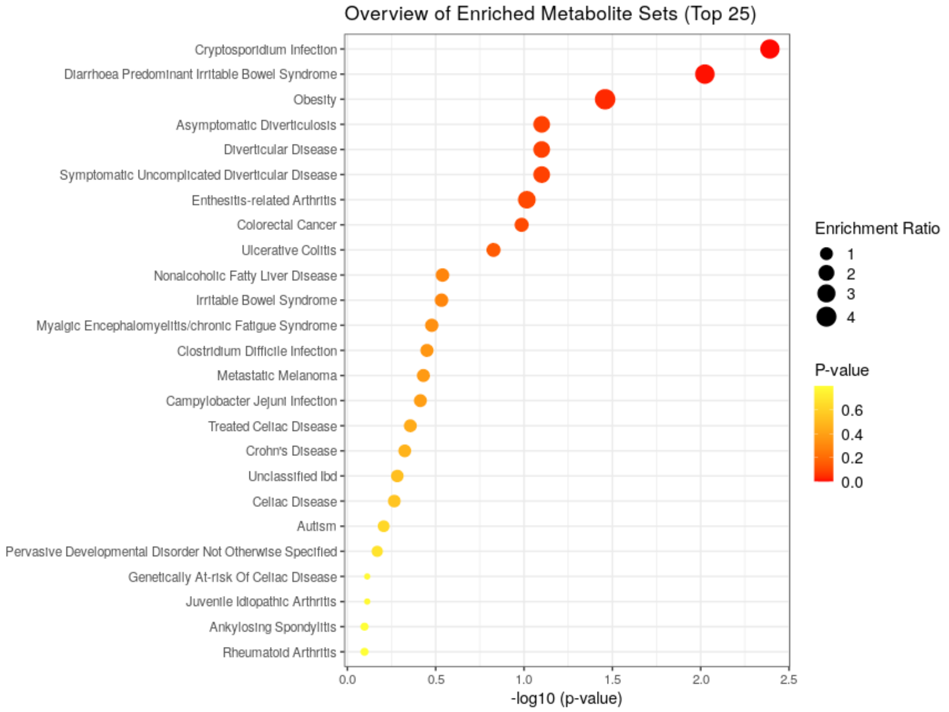


*Figure 27: Dot plot ranking disease features enriched in the human gut microbiota from two human faecal donors after exposure to 0,5% rhamnolipids for 7 days in the Simulator of the Human Intestinal Microbial Ecosystem (SHIME). Graph was obtained using the enrichment analysis tool on the MetaboAnalyst website.*

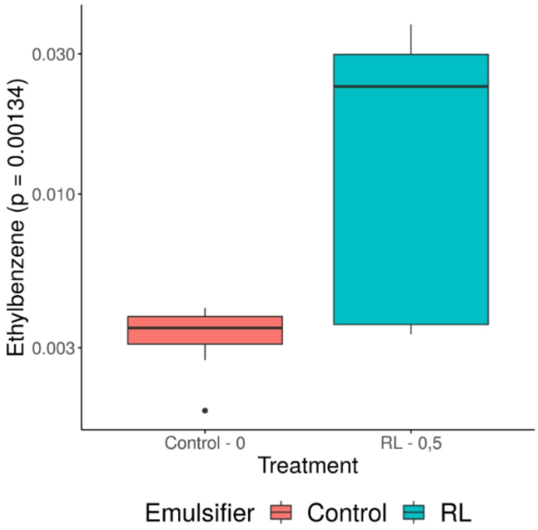


*Figure 28: Metabolites from the targeted metabolomics database annotated by the Enrichment Analysis tool on MetaboAnalyst as pointing towards Diarrhea dominated IBS for the treatment with 0,5% of rhamnolipids. Metabolites were detected in luminal suspensions from a 16 and 18 day SHIME experiment investigating the impact of TWEEN80 and rhamnolipids (0,05 m% and 0,5 m%) on the gut microbiota of two human faecal donors during a 7 day treatment period. Samples were from timepoint 2, 11, 12, 14 and 18 from SHIME 1 and timepoint 2, 9, 10, 12 and 16 from SHIME 2.*

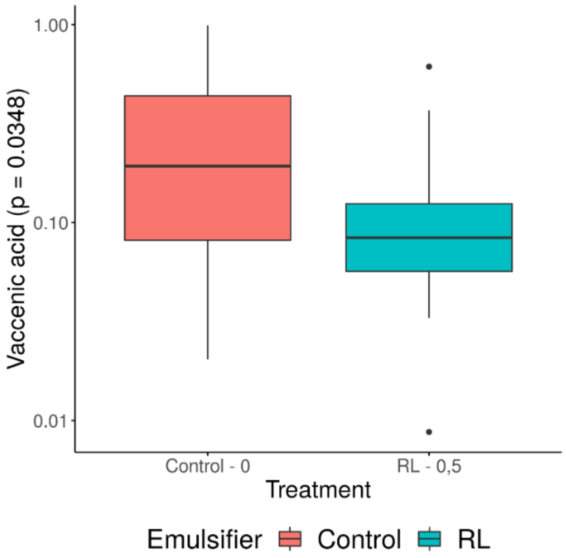


*Figure 29: Metabolites from the targeted metabolomics database annotated by the Enrichment Analysis tool on MetaboAnalyst as pointing towards Obesity for the treatment with 0,5% of rhamnolipids. Metabolites were detected in luminal suspensions from a 16 and 18 day SHIME experiment investigating the impact of TWEEN80 and rhamnolipids (0,05 m% and 0,5 m%) on the gut microbiota of two human faecal donors during a 7 day treatment period. Samples were from timepoint 2, 11, 12, 14 and 18 from SHIME 1 and timepoint 2, 9, 10, 12 and 16 from SHIME 2.*

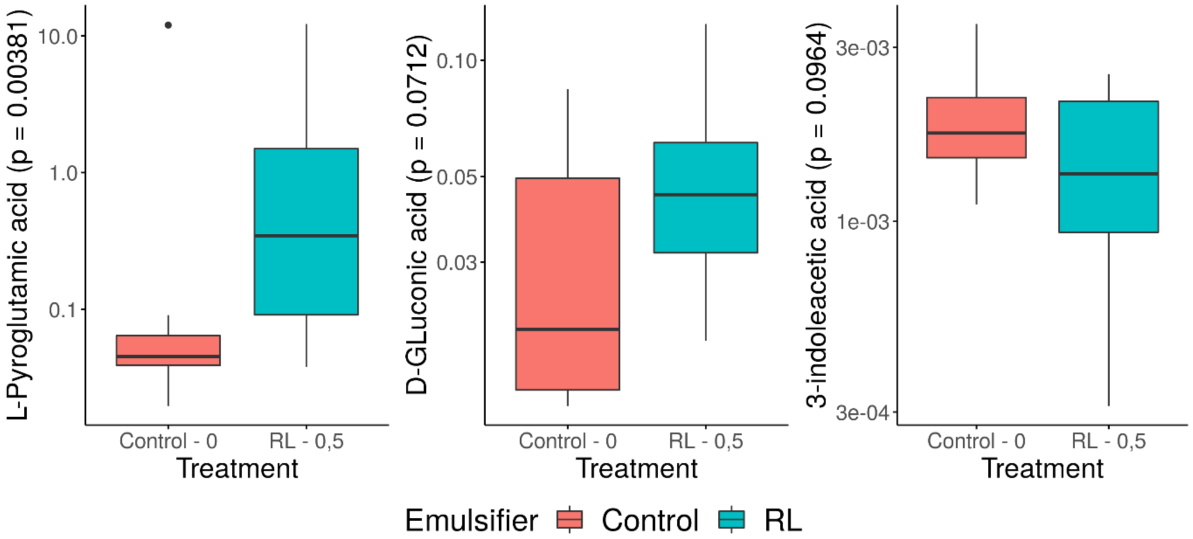


*Figure 30: Metabolites from the targeted metabolomics database annotated by the Enrichment Analysis tool on MetaboAnalyst as pointing towards Cryptosporium infection for the treatment with 0,5% of rhamnolipids. Metabolites were detected in luminal suspensions from a 16 and 18 day SHIME experiment investigating the impact of TWEEN80 and rhamnolipids (0,05 m% and 0,5 m%) on the gut microbiota of two human faecal donors during a 7 day treatment period. Samples were from timepoint 2, 11, 12, 14 and 18 from SHIME 1 and timepoint 2, 9, 10, 12 and 16 from SHIME 2.*

#### 4.2 Untargeted

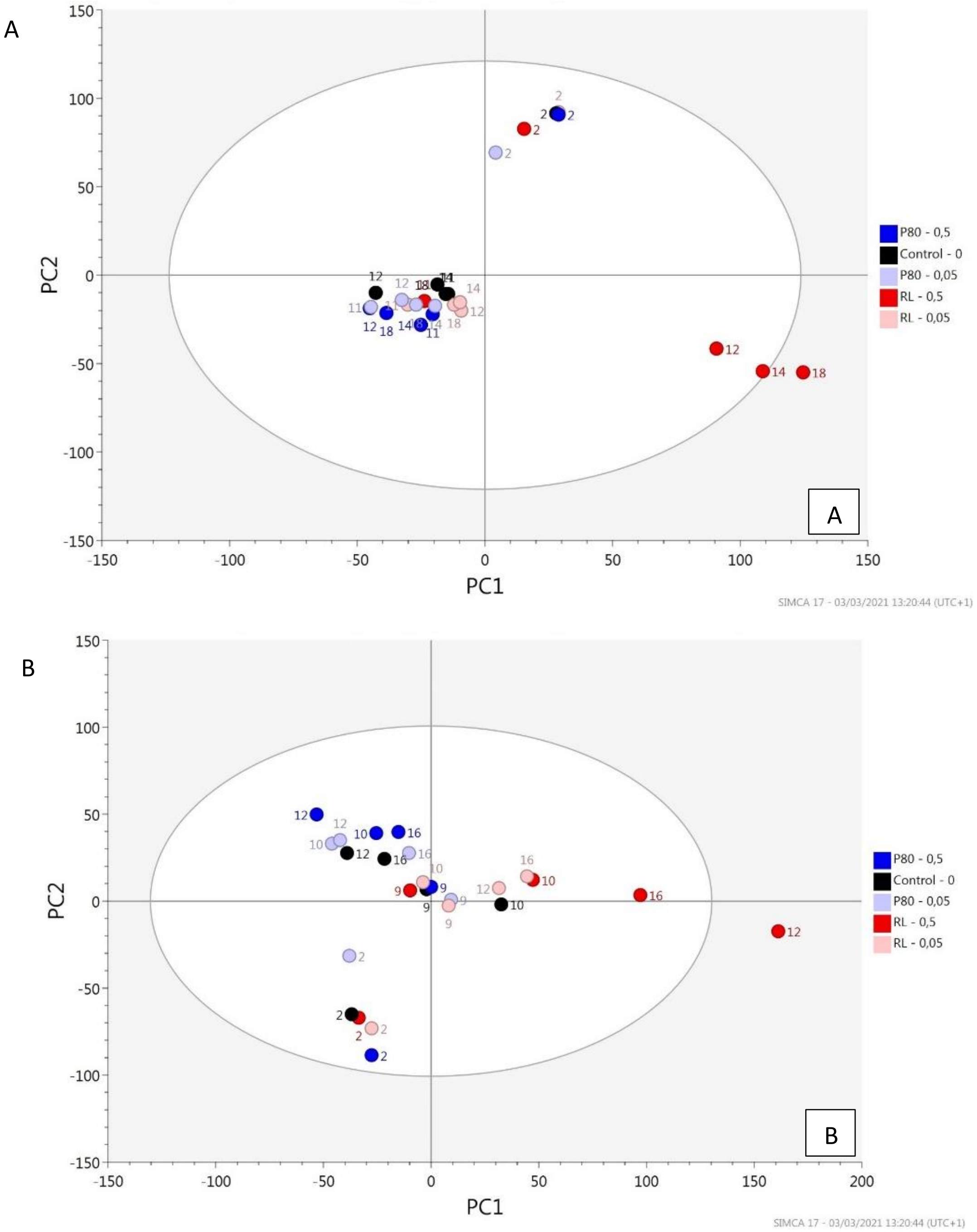


*Figure 31: Principle component analysis of untargeted metabolomics data extracted from luminal suspensions from a 16 and an 18 day SHIME experiment investigating the impact of TWEEN80 and rhamnolipids (0,05 m% and 0,5 m%) on the gut microbiota from two human faecal donors. Panel A = Donor 1, the donor selected for low emulsifier sensitivity. Panel B = Donor 2, the donor selected for high emulsifier sensitivity.*

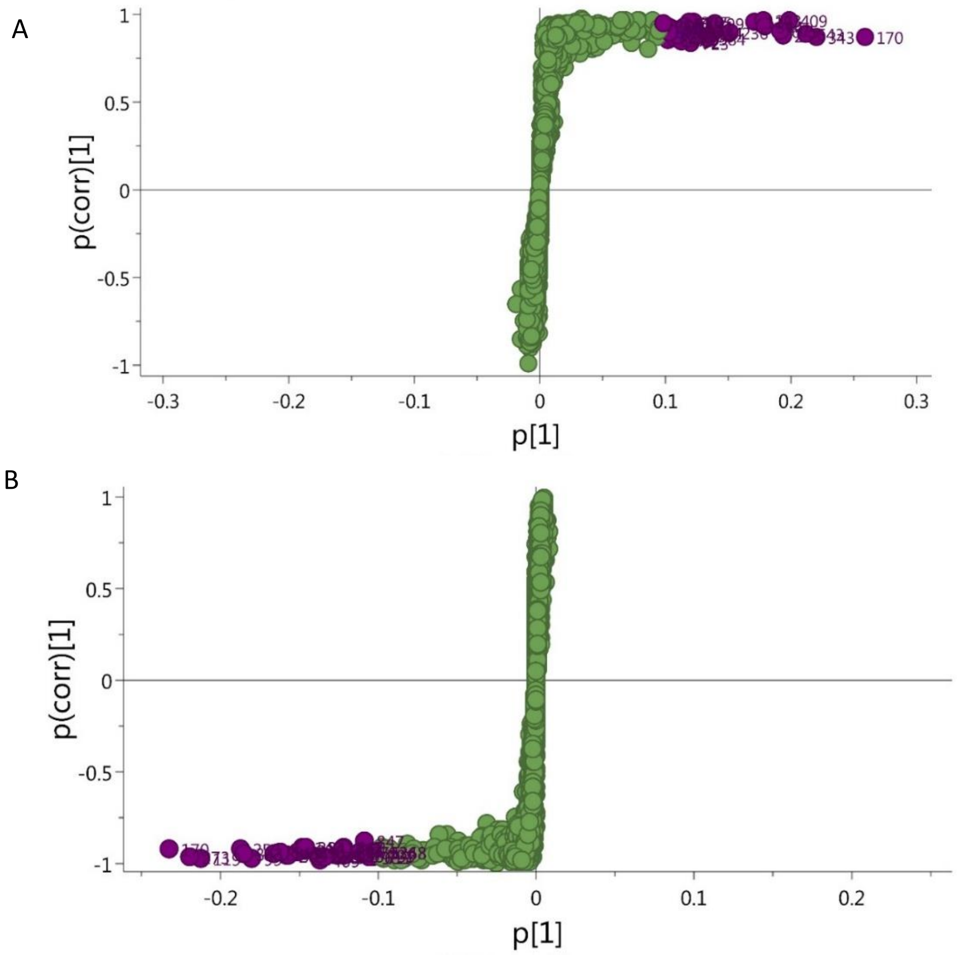


*Figure 32: S-plots obtained via orthogonal partial least squares analysis (OPLS-DA) on untargeted metabolomics data of luminal suspensions from a 16 and an 18 day SHIME experiment investigating the impact of TWEEN80 (0,05 m% and 0,5 m%) on the composition and functionality of the gut microbiota from two human faecal donors. The OPLS-DA models compared both soy lecithin treatments – 0,05 m%: A and 0,05 m% : B - with the control. P[1] represents the relative magnitude of the change. P(corr)[1] represents the confidence/reliability of the effect. As a result the data points in the upper right and lower left corners are the least likely the result of spurious correlations (T. Zhang et al., 2015). All data points designate certain metabolites. Metabolites were considered potential biomarkers (purple) when p[1] > 0,05 and p(corr)[1] > 0,4.*

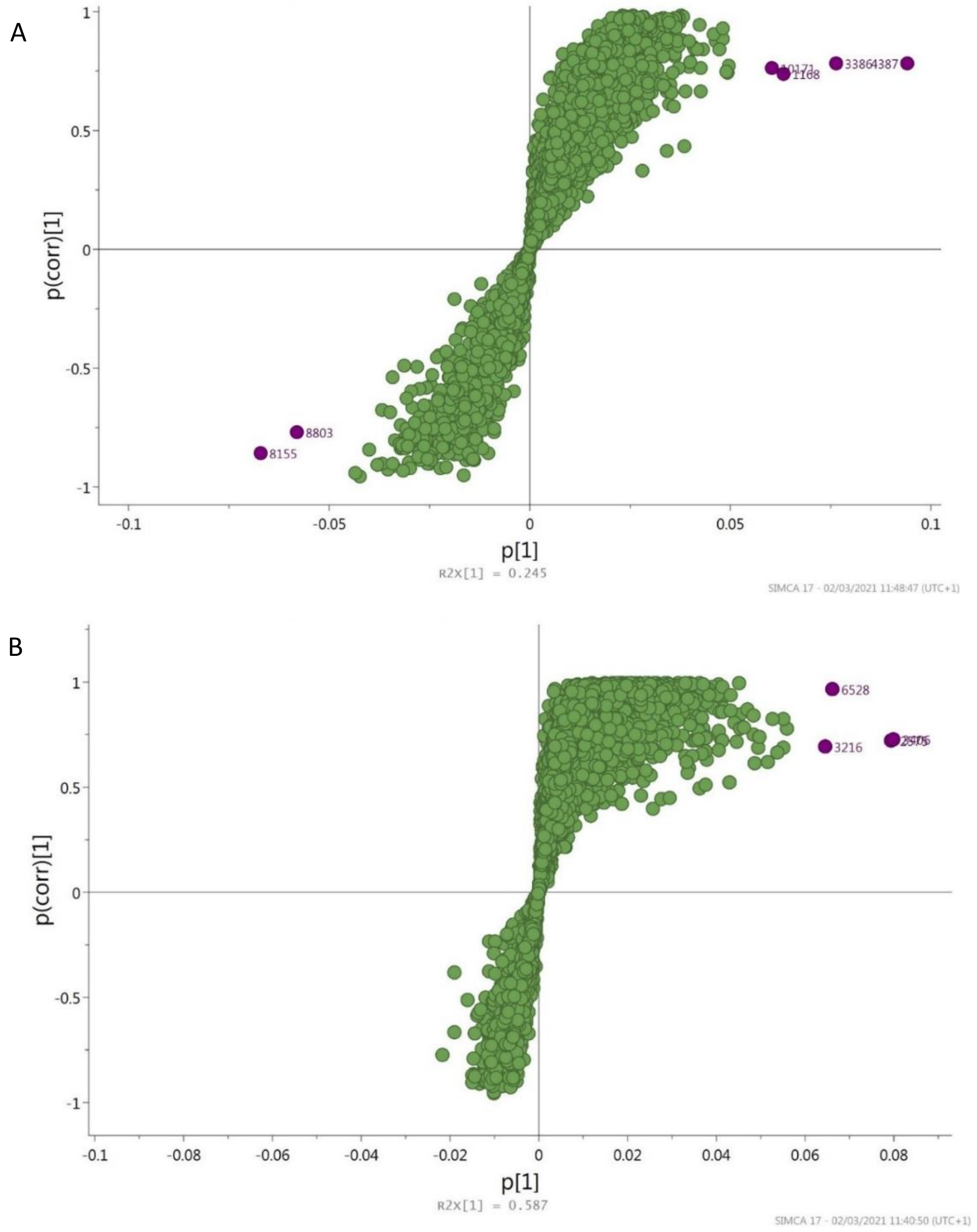


*Figure 33: S-plots obtained via orthogonal partial least squares analysis (OPLS-DA) on untargeted metabolomics data of luminal suspensions from a 16 and an 18 day SHIME experiment investigating the impact of rhamnolipids (0,05 m% and 0,5 m%) on the composition and functionality of the gut microbiota from two human faecal donors. The OPLS-DA models compared both soy lecithin treatments – 0,05 m%: A and 0,05 m% : B - with the control. P[1] represents the relative magnitude of the change. P(corr)[1] represents the confidence/reliability of the effect. As a result the data points in the upper right and lower left corners are the least likely the result of spurious correlations (T. Zhang et al., 2015). All data points designate certain metabolites. Metabolites were considered potential biomarkers (purple) when p[1] > 0,05 and p(corr)[1] > 0,4.*
